# Repeated exposure to viral and bacterial danger signals in Parkin deficient mice induces inflammation but fails to trigger parkinsonism

**DOI:** 10.64898/2026.09.22.753365

**Authors:** Amandine Even, Sriparna Mukherjee, Nicolas Giguère, Morgane Brouillard-Galipeau, Nathalia Luisa Oliveira, Priyabrata Halder, Claudie Beaulieu, Romain Cayrol, Nathalie Van Den Berge, Samantha Gruenheid, Louis-Eric Trudeau

## Abstract

Experimental models of Parkinson’s disease (PD) traditionally involve direct perturbations of the functioning of dopamine (DA) neurons. However, PD is increasingly recognized as multifactorial in origin and involving both cell-autonomous vulnerability factors and non-cell-autonomous triggers including inflammatory signals deriving from bacterial or viral infections. Notably, genes associated with early-onset familial PD as *PRKN* (Parkin) and *PINK1*, are ubiquitously expressed and have been implicated in immune regulation, further supporting a role for inflammation in disease etiology. In the present study, we tested the hypothesis that alternate Polyinosinic:polycytidylic acid (Poly(I:C)), mimicking viral infection, and lipopolysaccharides (LPS), mimicking bacterial infection, may induce sufficient brain inflammation to impair the DA system in Parkin-deficient mice and recapitulate some of the systemic features of PD. We developed a protocol with alternating Poly(I:C) and LPS, inducing acute peripheral inflammation in both genotypes, with consistent body weight loss and an increase of fecal lipocalin-2. Modest levels of neuroinflammation were present six months after the treatment, with higher levels of the microglial marker Iba1 in the dorsal striatum and reduced levels of the astrocyte marker GFAP in the mesencephalon, potentially indicating reduced neuronal support in the longer term. Finally, we find that this alternating Poly(I:C)/LPS treatment does not induce dopaminergic denervation or motor dysfunctions, even in Parkin KO mice. Taken together, our findings suggest that repeated short-term exposure to pathogen-derived signals is insufficient to induce DA system impairment in young adult Parkin deficient mice. Instead, induction of PD-like pathology may require a convergence of factors, including a prolonged or recurrent inflammatory burden, aging, genetic susceptibility, and potentially additional environmental or cellular stressors, that together create a pathogenic “perfect storm”. Thus, in its current form, the paradigm developed in the present study should not be regarded as a fully representative model of PD. Rather, it provides proof of concept that temporally alternating inflammatory stimuli can shape non-motor symptom-like phenotypes and underscores the need for future animal model research to incorporate interacting risk factors, disease duration, and age-dependent vulnerability.

## Introduction

The progressive degeneration of dopamine (DA) neurons within the substantia nigra (SN) in Parkinson’s Disease (PD) is a critical contributor to the characteristic motor symptoms of this disease such as bradykinesia, postural instability, rigidity of muscle tone and tremors (Tenchov et al., 2025). PD also includes a range of non-motor symptoms including gastrointestinal, olfactory and autonomic dysfunctions, sleep disturbances, dementia or depression, some of which appear years before the onset of motor symptoms and diagnosis (Palanivel et al., 2025).

Experimental PD models have conventionally targeted DA neuronal functions through direct perturbation. This includes DA depletion models using toxins like 6-OHDA (6-hydroxydopamine) or 1-methyl-4-phenyl-1,2,3,6-tetrahydropyridine (MPTP) (Dovonou et al., 2023). But these methods typically induce rapid and severe damage, that does not reflect the slow progressive nature of the disease. They also fail to capture the growing recognition that PD is in most cases a multifactorial disease involving cell-autonomous vulnerability factors and non-cell-autonomous triggers, including inflammatory signals deriving from bacterial or viral infections (Giguère et al., 2018; Khan et al., 2023; Tchung et al., 2026). Increasing evidence suggests that PD results from the convergence of multiple factors, including aging, genetic susceptibility (e.g. mutations in LRRK2, PRKN, PINK1, and SNCA) and environmental exposures. More comprehensive animal models are therefore needed to elucidate how genetic vulnerability interacts with peripheral signals to promote neuroinflammation and initiate disease onset.

It has been shown in multiple studies that people with PD exhibit signs of chronic inflammation (Harms et al., 2023; Tansey et al., 2022). Consistent with this, high concentrations of pro-inflammatory cytokines (Dzamko, 2023) and an alteration of immune cell populations in the blood have been observed (Pike et al., 2024). In the brain, activated microglia (Gerhard et al., 2006; McGeer et al., 1988; Tansey et al., 2022) and infiltrated T cells can be detected (Galiano-Landeira et al., 2020; Ma et al., 2025). While such inflammatory signals were previously considered as consequences of the disease, they are now increasingly investigated as potential contributors to its onset. Increasing evidence suggests that bacterial or viral exposure may play a role in the development of PD (Mercado et al., 2024; Smeyne et al., 2020). Indeed, multiple studies have linked viral infections (e.g., Influenza virus, Hepatitis C virus) to a higher prevalence of PD (Leta et al., 2022; Mercado et al., 2024; Smeyne et al., 2020). Similarly, epidemiological studies have highlighted that bacterial infection (e.g., *Helicobacter pylori* or *Mycobacterium tuberculosis*) elevates the risk of developing PD (Mercado et al., 2024; Shen et al., 2016, 2017; Smeyne et al., 2020).

Interestingly, some genetic mutations associated with an increased risk of the disease onset also appear to be linked to alterations in immune function. This is the case for loss-of-function mutations in *PRKN* or *PINK1*, major genetic contributors to early-onset forms of PD. Parkin and PINK1, proteins encoded by these genes, collaborate in the clearance of damaged mitochondria (Pickrell and Youle, 2015), also play a role in immune response regulation. Previous *in vitro* work revealed that PINK1 and Parkin play a role in repressing mitochondrial antigen presentation (MitAP) by antigen-presenting cells (Matheoud et al., 2016). Moreover, knock-down or knockout (KO) of *Pink1* or *PRKN* significantly impact microglial properties, exacerbating their transition to a reactive pro-inflammatory state following LPS or IFN-γ stimulation (Dionísio et al., 2019; Mouton-Liger et al., 2018; Sun et al., 2018). *In vivo*, while the absence of Parkin or PINK1 typically is not associated with overt neurodegeneration in mice (Paul and Pickrell, 2021), the induction of inflammatory stress triggers an intensified inflammatory and immune response (Kazanova et al., 2024; Mukherjee et al., 2026; Recinto et al., 2025) and can lead to motor dysfunctions (Matheoud et al., 2019). Recent studies have also shown that infection of PINK1 KO mice with the intestinal bacterium *Citrobacter rodentium* induces greater elevation of IL-17 cytokine and CXCL1 chemokine in the blood, accompanied by a microglial overactivation in PINK1 KO mice (Mukherjee et al., 2026). Furthermore, repeated infections with *Citrobacter rodentium* in a similar mouse model promote the establishment of a mitochondrial antigen-specific CD8^+^ T cell population (Matheoud et al., 2019). The subsequent migration of this cell population into the brain appears sufficient to induce degeneration of DA neurons and motor perturbations (Elemeery et al., 2024).

Lipopolysaccharide (LPS) is a bacterial endotoxin that can be administered to mimic a bacterial infection, while Polyinosinic:polycytidylic acid (Poly(I:C)) is a double-stranded RNA mimicking viral infection. Previous work has shown that a single intracerebral administration of a high dose of LPS or Poly(I:C) induces robust microglial activation and a rapid alteration of nigral DA neurons, characterized by a marked reduction in tyrosine hydroxylase (TH)-positive neurons (da Silva et al., 2024; Deleidi et al., 2010; Deng et al., 2020). Notably, intraperitoneal LPS injection also induces robust systemic inflammation and results in brain inflammation (da Silva et al., 2024). Prolonged exposure over several months can induce pronounced dopaminergic neuronal loss in the mesencephalon of Parkin-deficient mice(Frank-Cannon et al., 2008), and also leads to mild motor impairment in WT mice. Furthermore, peripheral LPS injection induces enteric inflammation in addition to neuroinflammation, mimicking the exacerbated intestinal inflammation observed in people with PD compared to healthy volunteers (Campagnolo et al., 2024; Dumitrescu et al., 2021). Together, these findings underscore the value of animal models that integrate genetic susceptibility with chronic inflammatory stress to better recapitulate the multifactorial nature of PD. However, models requiring the use of live bacteria typically require specialized facilities, while those relying on a single pathogen-derived signal often require repeated administrations over several months, making them less likely to of general use. Moreover, one potential issue with repeated administration of the same pathogen-derived signal is that it is likely to lead to immune system adaptations or tolerance (Kehl et al., 2004; Kim et al., 2024b), further limiting the magnitude and duration of the inflammatory response. Consequently, reproducing sustained and clinically relevant inflammation in rodents may require the integration of multiple distinct inflammatory insults to generate persistent immune activation and neuropathological changes that more closely resemble those observed in humans. Interestingly, the presence of a viral infection has been shown to facilitate a secondary bacterial infection and enhance the associated inflammatory response (McCullers, 2014; Sencio et al., 2020). Furthermore, people with PD seem to present a more diverse serology compared to healthy individuals (Bu et al., 2015), possibly resulting from previous exposure to bacteria as well as viruses.

Therefore, in the present study, we tested the hypothesis that alternate injections of Poly(I:C), mimicking viral infection, and LPS, mimicking bacterial infection, may induce sufficient brain inflammation to impair the DA system in Parkin-deficient mice. To characterize this model, we examined not only the effect on DA neurons but also on peripheral and central inflammation, as well as gut inflammation, aiming to establish a model that replicates several dysfunctions associated with PD in humans. We find that a dual exposure to Poly(I:C) and LPS induces peripheral inflammation and reduces the number of GFAP-positive astrocytes in Parkin KO mice at 6 months post exposure, suggesting the emergence of early alterations in astrocyte-neuron interactions. Although the dual inflammatory challenge paradigm used here did not induce detectable dopaminergic denervation 6 months post exposure, it generated a phenotype consistent with the early, non-motor stage of PD, characterized by persistent peripheral inflammation and subtle CNS alterations. This finding is consistent with the human disease trajectory, in which the prodromal phase may precede the onset of motor symptoms by two decades or longer. If validated through long-term studies, this model could provide a valuable platform for identifying early, subtype-specific biomarkers during the pre-motor phase, when therapeutic intervention may be most effective in modifying disease progression.

## Materials and methods

### Animals

All procedures involving animals were conducted in strict accordance with the Guidelines defined by the Canadian Council on Animal Care. The experimental protocols were approved by the animal ethics committee (CDEA) of the Université de Montréal. Mice were housed and maintained under reversed 12h light/dark cycles (so experiments were performed during the animals’ dark active phase), at constant temperature (21°C) and humidity (60%), with access to food and water *ad libitum*. Parkin KO mice (Itier et al., 2003) were backcrossed (>10 generations) with C57BL/6NCrl mice (Charle Rivers, Strain Code 027, RRID:IMSR_CRL:027). The genotyping was performed with a KAPA HotStart^®^ Mouse Genotyping Kit (Roche, Cat#KK7352), containing a KAPA2G Fast Hot Start DNA polymerase. The following primers were used to identify Parkin WT and KO mice: TGCTCTGGGGTTCGTC (WT, forward), TCCACTGGCAGAGTAAATGT (WT, reverse), CCTGCTTGCCGAATATCAT (KO-Neo, forward), AAGGCGATAGAAGGCGATG (KO-Neo, reverse).

### Systemic drug administration and animal monitoring

We previously showed that four gavages with C. rodentium were sufficient to induce the onset of motor dysfunctions in Pink1 KO mice (Matheoud et al., 2019), informing experimental design for this study. Young adult (8-12 weeks old) Parkin KO mice with their sex and age-matched wild-type (WT) littermates received intraperitoneal (i.p.) injections, once a week for four weeks. Depending on the model tested, mice received either 3mg/kg of LPS (from Escherichia coli O111:B4; Invivogen, Cat#tlrl-eblps), or 20mg/kg of Poly(I:C) (high molecular weight, Invivogen, Cat#tlrl-pic) or saline (as a vehicle control, Stevens, Cat#184-116,). The body weight of all mice was monitored before (time 0) and 3-5 days following each injection.

Three experimental paradigms were used to compare neuroinflammatory response to single vs. dual inflammatory challenge: (1) mice were injected four times with LPS (n=3-7 mice per group), (2) mice received four injections of Poly(I:C) (n=5-7 mice per group) or (3) mice received four injection, alternating Poly(I:C) or LPS (n=9-16 mice per group). For the Poly(I:C)/LPS model (**Suppl. Fig. 1A**), experiments were performed on two independent cohorts, one year apart, and the data were pooled for analysis. Four mice developed health issues unrelated to the experimental procedures months after completion of behavioral testing and were excluded only from subsequent analyses.

### Fecal lipocalin and calprotectin assays

Fecal samples were collected at predetermined time points following injection with saline or Poly(I:C)/LPS from WT and Parkin KO mice and placed into pre-weighed 1.5 mL microcentrifuge tubes. After measuring the weight, 10 μL of 0.1% Tween20 in PBS was added for every mg of feces. The samples were then incubated on ice for 15 min, followed by vigorous vortexing for 45 min at 4°C using a freezer sample box secured to a vortex. Next, the tubes were centrifuged at maximum speed for 15 min at 4°C, and the resulting supernatants were transferred to fresh microcentrifuge tubes before storage at -80°C.

To assess gut inflammation, lipocalin-2 levels were measured using the Mouse Lipocalin-2/NGAL DuoSet ELISA kit (R&D Systems, Cat#DY1857, RRID:AB_3678814) and Calprotectin levels were measured using the Mouse S100A8 / S100A9 Heterodimer Duo Set ELISA Kit (R&D Systems, Cat#DY8596-05). The protein standards, as well as the capture and detection antibodies, were reconstituted according to the manufacturer’s guidelines. The 1% blocking buffer was prepared by dissolving bovine serum albumin (Sigma-Aldrich, Cat#A4503-100G) in PBS, followed by vacuum filtration. A Nunc MaxiSorp™ 96-well plate (Invitrogen, Cat#44-2404-21) was coated with capture antibody and incubated covered overnight at room temperature. The plate was then washed three times using a wash buffer (0.05% Tween20 in PBS1X). To avoid non-specific binding, a blocking buffer was added and incubated for 1–2h before another washing step. Standards were prepared through serial 1:2 dilutions as instructed by the manufacturer, while fecal samples were diluted 1:10 in blocking buffer, added to the wells and incubated for 2h at room temperature before another washing step. Next, detection antibody was added to each well, followed by a 2h incubation at room temperature and additional washing. Each well received streptavidin-HRP before being incubated in the dark at room temperature for 20 min, followed by another wash cycle. For signal detection, a substrate solution combining Color Reagent A (Stabilized Peroxide Solution) and Color Reagent B (Stabilized Chromogen Solution) was added and incubated in the dark at room temperature for 15 min for Lipocalin-2 and 20 min for Calprotectin. To halt the reaction, 1 N H₂SO₄ stop solution was added to each well. Finally, absorbance was measured at 450 nm, with a correction reading at 620 nm for Lipocalin-2 and using a wavelength correction at 540 nm or 570 nm for Calprotectin.

### Analysis of fecal short chain fatty acids (SCFA)

Fecal samples were collected at specified time points after the injection of saline or Poly(I:C)/LPS in WT and Parkin KO mice. The SCFA analysis was performed by Microbiome Insights (Richmond, British Columbia, Canada). Samples were analyzed by GC-FID (Gas chromatography flame ionization detector) and the testing method was adapted from *Zhao et al* (Zhao et al., 2006). Briefly, feces samples were resuspended in MilliQ water and homogenized using an MP Bio FastPrep system for 1 min at a speed of 4 m/s. To achieve a final pH of 2, 5M HCl was added to acidify the fecal suspensions. Following acidification, the suspensions were incubated and centrifuged at 10 000 RPM to separate the supernatant. The fecal supernatants were then spiked with 2-ethylbutyric acid to reach a final concentration of 1 mM. The extracted SCFA supernatants were stored in 2 mL GC vials with glass inserts. A standard cocktail for acetic acid, butyric acid, propionic acid, isobutyric acid, isovaleric acid, valeric acid, hexanoic acid and heptanoic acid was prepared (4 dilutions for each) and was used as a reference for the detection of SCFAs in the samples. The detection was performed using a Trace 1310 gas chromatography analyzer (Thermo) equipped with a flame ionization detector.

### Behavioral analyses

The behavioral experiments were conducted four months after the last injection to allow for an incubation period during which chronic inflammation could act. After one week of habituation with the experimenter (10 min per day spent with each mouse), subjects were tested in the open field, the rotarod, and the pole test. The experiments were performed blindly, without knowledge of the genotype or treatment of the animals.

#### Open Field

Locomotor behavior was recorded using an infrared actimeter (Superflex sensor version 4.6, Omnitech) with the Fusion software (v5.6 Superflex Edition, RRID:SCR_017972). Subjects were tested directly for a total of 30 min in the chamber without prior acclimation. We focused on analyzing the following parameters: total movement time, total distance traveled, number of vertical episodes, time spent in the center, and time spent in the periphery.

#### Pole test

To assess their general motor functions, mice were tested on a pole test for 3 trials, separated by 10 min of rest. For each trial, the mouse was acclimated to the cage in which the pole was placed for approximately 20s. Subsequently, the animal was placed head-upward on top of a vertical rough-surfaced pole (diameter 1 cm; height 48 cm). The time until it descended to the floor was recorded, with a maximum duration of 120s. This time was divided into the time required to turn and the time necessary for the mouse to fully climb down. The first trial was considered a habituation, and the results are presented as the mean of the last two trials.

#### Rotarod

Motor coordination was evaluated with a rotarod apparatus (Harvard Apparatus, Cat#LE8205). Mice were pre-trained on the rotarod to reach a stable performance. On the experimental day, mice were placed on the rotarod, rotating at a constant speed of 4 rpm. They remained on the rotarod until either one min had elapsed, or for a maximum of 5 attempts at placing them on the rod. The tests started one day after the training and consisted of 3 sessions per day, with 10 min of rest between each session, repeated over 3 consecutive days. Mice were tested on the device with an accelerated rotation of 4 to 40 rpm, over a 6 min period or as soon as the mouse fell from the accelerating rod.

### Tissue collection

Tissues were collected six months after the last injection. Mice were weighed and then anesthetized with an i.p. injection of sodium pentobarbital. Fluids and tissues were then collected in the following order.

#### Blood collection and serum preparation

Blood was collected via cardiac puncture, kept 20 min at room temperature before being centrifuged at 2500g 15 min at 4°C to obtain serum. A 2-fold dilution of serum was prepared with PBS pH∼7.5 and kept at -80°C before analysis.

### Spleen collection and spleen index

The spleens were carefully collected and weighed. The spleen index was calculated as spleen mass (mg)/mouse body mass (g).

#### Colon collection and colon index

Colons were sectioned between the cecum and the anus, feces were carefully removed and tissues were weighed. The colon index was calculated as the colon mass (mg)/mouse body mass (g). Then colons were washed with PBS to remove all fecal contents, sectioned in distal and proximal portions and placed at 4°C in 4% PFA until embedding.

#### Brain collection

Mice were perfused through the heart with cold PBS followed by 4% PFA. Two days after, brain samples were placed in a 30% sucrose solution for 48h to facilitate cryoprotection and then frozen. Brain samples were frozen using dry ice and stored in -80°C until sectioning.

#### Stellate ganglion collection

Left and right stellate ganglia, located laterally to the longus colli muscles and across the second rib, were removed, as described by *Scherschel et al* (Scherschel et al., 2020). They were stored at 4°C in 4% PFA until paraffin embedding.

### Quantification of serum cytokines and chemokines by multiplex immunoassay array

Serum cytokine and chemokine concentrations were quantified by Eve Technologies (Calgary, Canada) using a 32-plex (MD32 Mouse Cytokine/Chemokine 32-Plex Discovery Assay® Array, Eve technologies) immunoassay including: Eotaxin, G-CSF, GM-CSF, IFNγ, IL-1α, IL-1β, IL-2, IL-3, IL-4, IL-5, IL-6, IL-7, IL-9, IL-10, IL-12p40, IL-12p70, IL-13, IL-15, IL-17A, IP-10, KC, LIF, LIX, MCP-1, M-CSF, MIG, MIP-1α, MIP-1β, MIP-2, RANTES, TNFα and VEGF-A.

### Embedding and immunohistochemistry of the colon and stellate ganglion

Paraffin embeddings were performed by the Histology Core Facility of the Institute of Research in Immunology and Cancerology (IRIC) at the Université de Montréal. Briefly, fixed colons were dehydrated by using graded ethanol, cleared and embedded in paraffin wax.

#### Colon immunohistopathology

Distal and proximal portions were serially sectioned at 4μm, mounted onto glass slides, air-dried, and stained with hematoxylin and eosin (H&E). Briefly, sections were cleared in three changes of xylene (3 min each), rehydrated through graded ethanol (three changes of 100% ethanol and one change of 95% ethanol, 3 min each). Then sections were rinsed in distilled water. Sections were stained with hematoxylin (Epredia, Cat#7221) for 20s, rinsed under running tap water for 1 min, immersed in bluing reagent (Epredia, Cat#7301) for 1 min, rinsed again for 1 min, and counterstained with eosin Y (Epredia, Cat#7111) for 10s. Sections were subsequently dehydrated in 95% ethanol for 2 min followed by two 2-min washes in 100% ethanol. Then sections were cleared in xylene with two 2-min washes, and coverslipped using Permount mounting medium (Fisher Scientific, Cat#SP15). Histopathological scoring was conducted blindly by an expert board-certified pathologist based on the scoring criteria of colon lesions: inflammatory cell type (significant presence of eosinophils, lymphocytes, macrophages, neutrophils); inflammatory cell infiltration (0-4); inflammatory cell localization (depth) (0-4); submucosal edema (0-4); surface epithelial injury (0-4); mucosal necrosis (0-4); goblet cell/enterocyte ratio decrease (0-4); gland loss (0-4). Slides were scanned on Aperio Scanscope (Leica, RRID:SCR_018457) and visualized on NDP.view2 software (Hamamatsu Photonics, RRID:SCR_025177).

#### Stellate ganglion immunohistochemistry

Briefly, the paraffin-embedded tissue was cut into 4-µm-thick sections using a Rotary microtome (Leica RM2235, RRID:SCR_026059) and mounted on SuperFrost Plus glass adhesion slides (Thermo Scientific, Cat#22034980). Tissue sections were deparaffinized in xylene and rehydrated in a series of alcohol baths. The sections were stained using the Benchmark Ultra autostainer (Roche Ventana, RRID:SCR_025506). The automated staining protocol included 5 steps. First a peroxidase-blocking was performed during 5 min, then an antigen retrieval was carried out using Cell Conditioner solution (CC1, pH8.5) for 4 min at 95°C. Primary tyrosine hydroxylase (TH, 1:10000, Abcam, Cat#ab112, RRID:AB_297840) and choline acetyltransferase (ChAT, 1:15000, Abcam, Cat#ab178850, RRID:AB_2721842) antibodies were incubated for 16 min at 37°C. Secondary antibody detection was performed using the ultraView Universal DAB Detection Kit (Roche), following by a bluing for 4 min. After staining, slides were washed in water with dishwashing detergent and dehydrated through graded alcohol baths. After clearing in xylene, slides were mounted and sealed using Eukitt (Merck) mounting medium and coversliped.

Images were acquired using a VS120 automated slide-scanner (Olympus, RRID:SCR_018411). The drawing tool from Aiforia (Aiforia cloud, RRID:SCR_022739) was used to delineate the boundaries of each section where positive staining needed to be quantified. Aiforia’s built-in analysis tool generated quantitative data on TH- and ChAT-positive staining of the stellate ganglia. A weighted sum of the generated RGB values was used to convert to grayscale intensity (0.299R + 0.587G + 0.114B). To calculate inverted optical density from grayscale intensity, the following formula was applied -log10(grayscale intensity/255).

### Immunohistochemistry on brain slices

Using a Leica CM1950 cryostat (Leica, RRID:SCR_018061), serial coronal sections (40µm thick) were cut at -21°C and collected in a cold cryoprotectant solution (30% ethylene glycol, 30% glycerol in 0.1 M phosphate buffer). The floating sections were washed three times (10 min each) with PBS. Permeabilization and blocking were performed under agitation for 60 min in a solution containing 0.3% Triton X-100, 5% goat serum, and 10% bovine serum albumin. For immunostaining, slices were incubated overnight under gentle agitation at RT with primary antibodies rabbit anti-TH (1:1000, Millipore Sigma, Cat#AB152, RRID:AB_390204), rat anti-DAT (1:1000, Millipore Sigma, Cat#MAB369, RRID:AB_2190413), guinea pig anti-Iba1 (1:1000, Synaptic System, Cat#234 308, RRID:AB_2924932) or chicken anti-GFAP (1:2000, Abcam, ab4674, RRID:AB_304558). Primary antibodies were subsequently detected with goat anti-rabbit AlexaFluor-488-conjugated (1:1000, Invitrogen, Cat#11008, RRID:AB_143165), goat anti-rabbit AlexaFluor-546-conjugated (1:1000, Invitrogen, Cat#11010, RRID:AB_2534077), goat anti-rat AlexaFluor-647-conjugated (1:1000, Invitrogen, Cat#21247, RRID:AB_141778), goat anti-guinea pig AlexaFluor-488-conjugated (1:1000, Invitrogen, Cat#11073, RRID:AB_2534117), or goat anti-chicken AlexaFluor-647-conjugated (1:1000, Biotium, Cat#20044, RRID:AB_10853617) secondary antibodies. For this, the sections were washed with PBS before being incubated for 2h at room temperature with secondary antibodies and DAPI (1:1000, Cat# D9542-1MG). After three additional PBS washes (10 min each), the slices were mounted onto Surgipath X-tra charged microscope slides (Leica, Cat#3800200), sealed with Fluoromount-G® mounting medium (Southern Biotechnology, Cat# 0100-01) and glass coverslips. and stored at 4°C until imaging.

### Confocal imaging

Immunohistofluorescence images were obtained using an Eclipse Ti2 inverted confocal microscope (Nikon, RRID:SCR_021068). To minimize nonspecific signal bleed-through, images were acquired sequentially using laser excitation 405, 488, 546 and 647nm. Three coronal sections from the striatum (AP +1.18, +0.14, and −0.94 mm relative to bregma) and three from the mesencephalon (AP -3,08, -3,28, and -3,52 mm relative to bregma) were selected from each brain for analysis **(Suppl. Fig. 1B)**. For TH and DAT quantification, large images were acquired with a 20X objective using the JOBS acquisition module. For quantification of microglia, 24 striatal and 14 mesencephalic sections were selected. Z-stacks were obtained using a 40x objective, with steps of 1µm.

### Image analysis

All the acquired images were processed using Nikon Imaging Software (NIS) Element software (version 6.20.02, Nikon, RRID:SCR_014329) with the Advance Research (AR) and NIS.ai modules or using Image J (RRID:SCR_003070). Custom scripts developed within NIS to facilitate image analysis are available at: https://github.com/Louis-EricTrudeau/Trudeau-lab/tree/main/Even-2026.

#### Quantification of TH/DAT fiber intensity and area

Regions of interest (ROIs) within the dorsal and ventral striatum were delineated for further analysis. Within the NIS software, image deconvolution using the Batch Deconvolution tool was performed prior to quantification to enhance image quality and reduce background noise. The rolling ball average and thresholding functions were applied to correct uneven background distribution. Mean values for fluorescence intensity and fiber area were calculated for the dorsal and ventral striatal ROIs and subsequently analyzed across experimental groups.

#### Microglial counting

Images were processed and analyzed using NIS software. Briefly, an extended depth of field (EDF) reconstruction of image stacks was performed, followed by an AI denoising **(Suppl. Fig. 1C)**. Microglia cells bodies were detected and counted using machine learning developed in NIS software. In mesencephalic slices, the TH area was selected to exclude regions peripheral to the DA cell body areas by setting intensity thresholds, determined based on several test images.

#### Volume and intensity of Iba1-positive microglia and GFAP-positive reactive astrocytes

The analysis was performed on a 3D reconstruction of image stacks with NIS software **(Suppl. Fig. 1D)**. Denoise and local contrast functions were applied to correct background and homogenize the intensity between samples. In the mesencephalon, only the region containing TH-positive neurons was selected for further analysis. DAPI^+^, Iba1^+^ or GFAP^+^ objects were identified by setting intensity thresholds and segmentation, determined based on several test images. The sum of the volume of Iba1^+^ and GFAP^+^ objects were calculated. Then, the mean fluorescence intensity of untouched Iba1 and GFAP signals was quantified across the measurement volume. The number of DAPI^+^ objects inside the GFAP volume was counted as number of GFAP-positive cells.

#### Microglia morphological analysis

Microglial morphological analysis was performed using ImageJ **(Suppl. Fig. 1E)**. The channel corresponding to the Iba1 marker, considered as microglia, was extracted from the entire Z-stack. The denoised image stacks were then merged to generate a 2D image while retaining partial 3D information (by preserving the maximum intensity of each pixel in the Z-stack). For the mesencephalon, only the Iba1 area that colocalized with the TH area was included in the analysis. Then, we used the following protocol described and developed by Cierna et al (Kim et al., 2024a). Several test images were used to define a fluorescence intensity threshold for distinguishing microglia, utilizing the MicrogliaMorphology and Biovoxxel ToolBox plugins (Jan Brocher, biovoxxel/BioVoxxel-Toolbox: BioVoxxel Toolbox v2.6.0. Zenodo; 2023). Minimum and maximum size parameters were then set to filter particles and exclude overlapping microglia using the MicrogliaMorphology plugin. Each 2D image was subsequently analyzed by the program: each identified cell was selected, assigned a unique identification number, and saved as an individual file. A skeleton representation was generated for each identified cell using the Skeletonize 2D/3D plugin, and skeleton parameters such as the number and length of processes were measured. Finally, a fractal analysis was performed for all identified cells using the FracLac plugin (Karperien, A., FracLac for ImageJ. Introduction.htm. 1999-2013) to determine various morphological parameters, including circularity, area, perimeter, and pixel count. Finally, the clustering of microglia into 4 groups (ramified, rod-like, ameboid, and hypertrophic) and the calculation of the percentage for each population were performed using RStudio.

### Statistical analysis

Data was analyzed with GraphPad Prism 10 software. Two-way ANOVA, three-way ANOVA or mixed-effect model (REML) were performed as appropriate. All statistical analysis results are reported in the **Supplementary Tables 1-12**. These global analyses were followed by Šídák’s multiple comparisons test. When necessary, logarithmic or square-root transformations were performed to improve normality and stabilize variances prior to applying the appropriate statistical tests. When significant interactions were observed, simple effects analyses were performed by fixing one factor and examining the effects of the remaining factors. When extreme values affecting the mean or skewed data distributions were detected, non-parametric rank-based tests (e.g., Mann–Whitney test) were used. Bonferroni correction was applied to adjust for multiple comparisons. Due to the high frequency of zero values in score data, additional analyses based on score presence or absence were performed using contingency tables and Fisher’s exact test. When significant, pairwise Fisher’s exact tests with Bonferroni correction were conducted. Graphs were generated using GraphPad Prism 10 and depict all data as mean ± SD.

## Results

### Repeated systemic administration of LPS but not of Poly(I:C) induces a tolerance revealed by a gradually attenuated body weight loss

Previous attempts to model aspects of PD in mice using peripheral LPS administration implicated protocols with multiple injections. In some cases, this was as high as two injections per week for 6 months (Frank-Cannon et al., 2008). Other studies used a single LPS injection followed by a prolonged observation period of 7 to 10 months (Qin et al., 2007). Both approaches can be considered impractical, perhaps explaining why they are rarely used. In a previous study, we observed that four gavages with *Citrobacter rodentium* were sufficient to induce the onset of motor dysfunctions in PINK1 KO mice (Matheoud et al., 2019). Here we tested the hypothesis that four i.p. injections of pathogen-associated molecular pattern (PAMP) are sufficient to trigger motor dysfunctions and perturbations of the dopaminergic system. In an initial experiment, we injected 2-month-old Parkin WT and KO mice i.p. with LPS or Poly(I:C), once a week for four weeks **(Fig. 1A)**. The weight loss profile exhibited four distinct phases, each occurring after LPS injection and accompanied by marked body weight loss. An overall change in body weight was observed in LPS-treated mice over time *(3-way ANOVA, treatment and time effects p<0.0001,* **Supplementary Table 1**) **(Fig. 1B)**. However, the extent of body weight loss decreased gradually with each injection and was insignificant by the fourth injection *(3-way ANOVA, treatment and time effect p<0.0001 for phase 1 to 3 and p=0.4303 for phase 4,* **Supplementary Table 2**), indicating tolerance to LPS. More specifically, body weight was decreased by 10.38% (±3.35) at the first peak of weight loss, followed by only 5.05% (±1.56), 4.93% (±1.37) and 3.11% (±3.89) across successive injections in the WT LPS treated group. Similar profiles were observed in Parkin KO mice, with a decrease of 13.09% (±3.18) after the first injection, followed by only 5.05% (±2.62), 3.91% (±1.85), and 3.75% (±3.62) in response to the subsequent injections. These observations are in line with previous work showing that repeated activation of the TLR4 receptor by LPS induces a tolerance phenomenon (Kehl et al., 2004; Liu et al., 2008; Musaelyan et al., 2018; Y. Yang et al., 2022).

**Figure 1:**
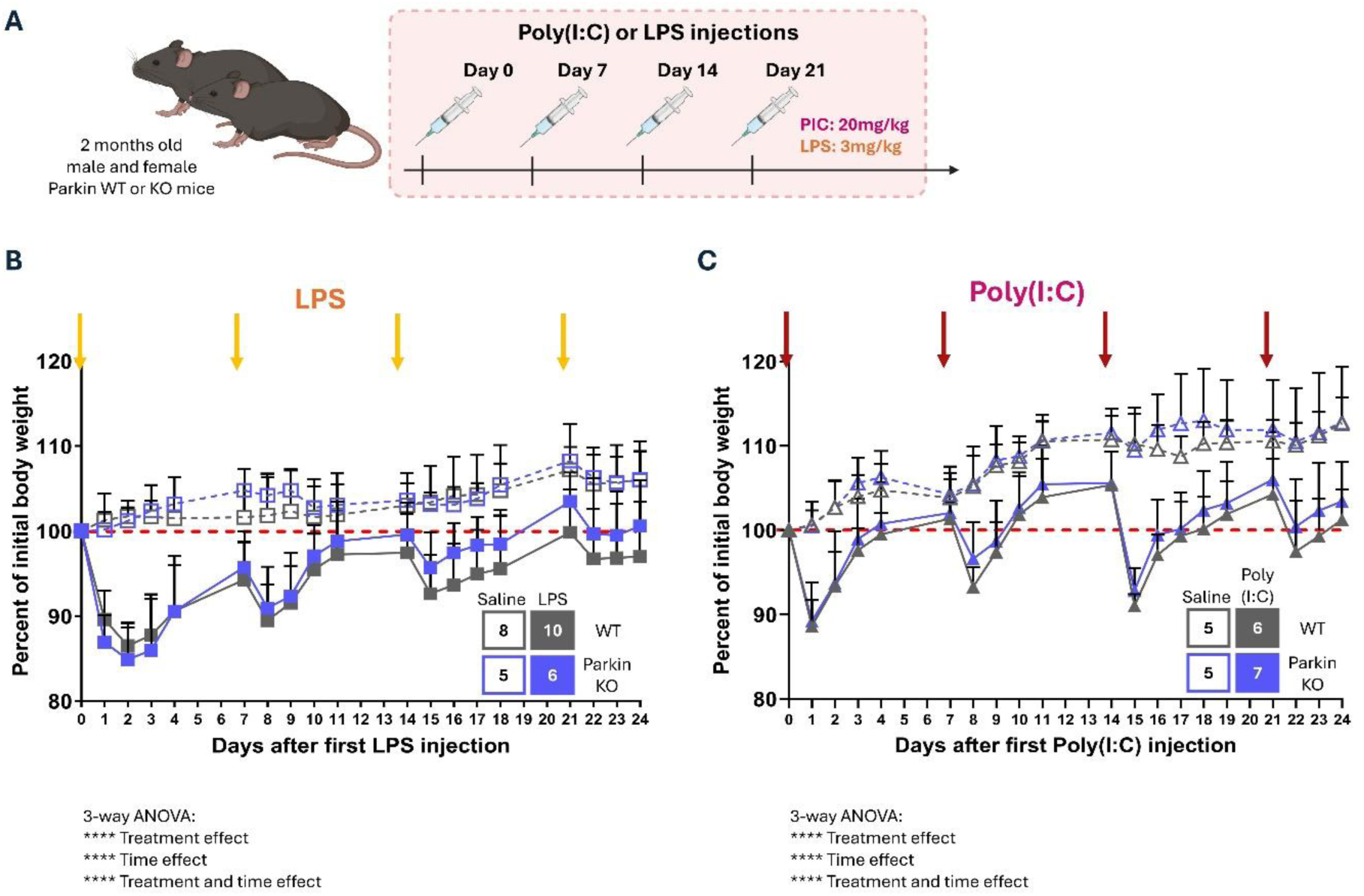
Repeated systemic administration of LPS but not of Poly(I:C) induces adaptations revealed by a gradually reduced body weight loss. **(A)** Schematic illustration of experimental design. Two-month-old male and female WT or Parkin KO mice received intraperitoneal injections of LPS (3 mg/kg) or Poly(I:C) (20 mg/kg), once weekly for four consecutive weeks. B-C, Body weight was measured before and for 3–5 days following each injection and represented in percentage of initial body weight. **(B)** LPS induced a marked reduction in body weight after the first administration, with progressively attenuated effects following subsequent injections, irrespective of genotype. n=8 for WT Saline, n=10 for WT LPS, n=5 for Parkin KO Saline and n=6 for Parkin KO LPS. **(C)** Poly(I:C) produced a reproducible decrease in body weight after each injection, although the magnitude of the effect was inconsistent over time. The overall pattern of body weight loss is preserved in KO mice. n=5 for WT Saline, n=6 for WT Poly(I:C), n=5 for Parkin KO Saline and n=7 for Parkin KO Poly(I:C). **B-C**, WT mice are shown in grey and Parkin KO mice in blue. Dotted lines represent saline-treated controls, and solid lines represent treated animals. Squares and triangles with error bars depict the mean ± SD per group of mice. Arrows indicate days of injection. Statistical significance was assessed using 3-way ANOVA (* p<=0.05, **** p<0.0001), the results of which are presented in **Supplementary Table 1**.

A second cohort of mice were injected four times with Poly(I:C). This PAMP is often used to model viral infections, and considered a potential contributing drivers of PD and other neurodegenerative diseases. However, no previous study has tested whether repeated injections of Poly(I:C) can be used to model PD. Like LPS, we found that Poly(I:C) impacted body weight over time *(3-way ANOVA, treatment and time effect p<0.0001,* **Supplementary Table 1**) **(Fig. 1C)**. Interestingly, less tolerance was observed after administration of this PAMP. Separate analysis of each post-injection period revealed a significant treatment and time interaction following all four Poly(I:C) injections *(3-way ANOVA, treatment and time effect p<0.0001 for phase 1 to 4,* **Supplementary Table 3**), suggesting a more sustained impact of Poly(I:C) compared to LPS. Furthermore, Poly(I:C), appears to have a more pronounced effect on body weight compared to LPS, although the first and third injections appeared to induce larger effects compared to the second and fourth. In WT mice, we observed a decrease of body weight of 11.36% (±3.08), 7.97% (±2.56), 13.52% (±3.46) and 6.72% (±4.42) in response to the four injections. A comparable effect occurred in Parkin KO mice, with decreases of 10.80% (±4.64), 5.25% (±3.70), 12.04% (±2.80) and 5.23% (±3.43) in response to the four injections.

These data indicate that repeated administration of PAMPs that mimic bacterial and viral infections do not induce the same adaptations over time. Furthermore, we found that loss of Parkin did not affect body weight loss induced by these PAMPs.

### Repeated systemic administration of Poly(I:C)/LPS induces similar acute peripheral inflammation in WT and Parkin KO mice

To overcome the tolerance observed in single-PAMP models, we alternated Poly(I:C) and LPS injections to establish a dual-PAMP inflammatory model **(Fig. 2A).** With this protocol, a persistent decrease in body weight was observed following all four injections (**Fig. 2B**). An overall interaction effect between time, treatment and genotype was observed (*3-way ANOVA, time, treatment and genotype p=0.0226,* **Supplementary Table 1**). When the analysis was stratified by injection phase, this interaction was only maintained during phase 3 (*3-way ANOVA, time, treatment and genotype p=0.0020,* **Supplementary Table 4**), whereas no significant interaction with genotype was observed in the other phases. These results suggest that the contribution of genotype to the interaction observed in the full model is phase-dependent and is primarily driven by phase 3. However, when data were analyzed at individual time points to fix this variable, no significant effect of the genotype was observed at days 14, 15, 16 and 17, indicating that its influence is distributed across the temporal trajectory rather than at specific time points. Otherwise, as expected, the Poly(I:C)/LPS model appears to induce a significant decrease in body weight after each injection and consistent over time (*3-way ANOVA, treatment and time effect p<0.0001 for phase 1 to 4,* **Supplementary Table 4**).

**Figure 2:**
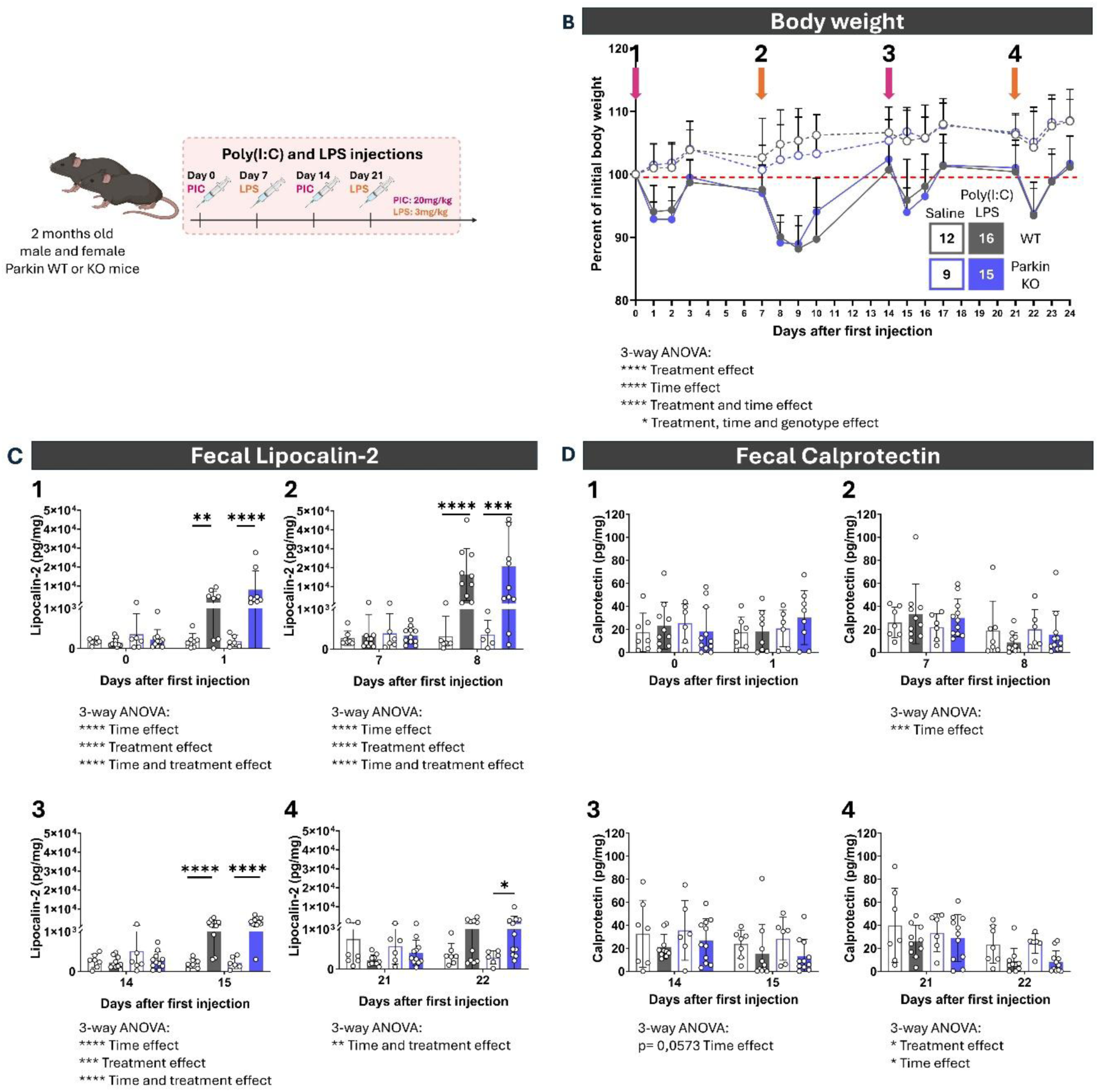
Repeated systemic administration of Poly(I:C)/LPS induces similar acute peripheral inflammation in WT and Parkin KO mice. **(A)** Schematic illustration of experimental design. Two-month-old male and female WT or Parkin KO mice received intraperitoneal injections of Poly(I:C) (20 mg/kg) and LPS (3 mg/kg) in alternation for four consecutive weeks. **(B)** The body weight was measured before and for 3 days following each injection and is represented by the percentage of initial body weight. Treated mice present a significant reduction after each injection, irrespective of the genotype. Dots with error bars depict the mean ± SD for each group. Dotted lines represent saline-treated controls, and solid lines represent treated animals. The numbered arrows from 1 to 4 represent the injection sequences, found in the following sub-figures. Statistical significance was assessed using 3-way ANOVA (* p<=0.05, **** p<0.0001), the results of which are presented in **Supplementary Table 1**. **C-D,** Feces were collected before and the day after each injection to measure **(D)** lipocalin-2 and **(E)** calprotectin by ELISA. The levels of fecal lipocalin-2 are increased after the first three injections, particularly in KO mice. A three-way ANOVA was first performed to assess the overall effect of each variable (** p<0.01, *** p<0.001, **** p<0.0001), the results of which are presented in **Supplementary Table 6**. When significant interactions were detected, the time factor was held constant and a two-way ANOVA followed by Šídák’s multiple-comparisons post hoc test was conducted. Significant comparisons are indicated above the bars (* adjusted p<=0.05, *** adjusted p<0.001, **** adjusted p<0.0001), the results of which are presented in **Supplementary Table 7**. Transformation of the data before testing has been done when necessary. However, the levels of fecal calprotectin appear stable. For panels **C-D,** bars represent the mean ± SD of the group and dots represent individual animals. Grey and blue are used respectively for WT and Parkin KO mice. Bars corresponding to saline are unfilled and outlined, while bars corresponding to Poly(I:C)/LPS are filled. n=12 for WT Saline, n=16 for WT Poly(I:C)/LPS, n=9 for Parkin KO Saline and n=15 for Parkin KO Poly(I:C)/LPS, from 2 independent cohorts.

Systemic injection of PAMPs is expected to induce peripheral inflammation, particularly in tissues such as the gut. Previous work revealed that people with PD have exacerbated intestinal inflammation compared to healthy volunteers (Campagnolo et al., 2024; Dumitrescu et al., 2021). Here, we measured fecal lipocalin-2 levels, a well-established biomarker of both acute and chronic intestinal inflammation (Chassaing et al., 2012; Thorsvik et al., 2017). Overall, we observed a significant increase in the levels of lipocalin-2 with treatment and time in both WT and Parkin KO mice (*3-way ANOVA on log transformed data, time and treatment effect, phase 1 to 3 p<0.0001, phase 4 p=0.0013,* **Supplementary Table 5**) **(Fig. 2C)**. When time points were analyzed separately, it appeared that significant differences occurred between the untreated and the PAMP-treated mice, specifically the day after the injections, (*2-way ANOVA on log transformed data, day 1 p=<0.0001, day 8 p<0.0001, day 15 p<0.0001, day 22 p=0.0044,* **Supplementary Table 6**). A comparison across genotypes revealed that fecal levels of lipocalin-2 were increased the day after the first three injections in WT mice (*2-way ANOVA on log transformed data, day 1 p=0.0448, day 8 p<0.0001, day 15 p<0.0001,* **Supplementary Table 6**) and after all injections in Parkin KO mice (*2-way ANOVA on log transformed data, day 1 p=0.0218, day 8 p=0.0005, day 15 p<0.0001, day 22 p=0.0198,* **Supplementary Table 6**). Calprotectin, a protein secreted by neutrophils during inflammation, is also elevated in the feces of people with PD (Aho et al., 2021; Dumitrescu et al., 2021; Hor et al., 2022). Here we observed that Poly(I:C) and LPS injections did not significantly alter the levels of fecal calprotectin in WT and Parkin KO mice **(Fig. 2D)**. To the contrary, fecal levels of calprotectin were reduced at day 8 and 22 compared to their baseline (*3-way ANOVA on log transformed data, time effect respectively p=0.0002 and p=0.0150,* **Supplementary Table 7**), with a tendency to decrease also at day 15 (*3-way ANOVA on log transformed data, time effect, p=0.0573,* **Supplementary Table 7**). Notably, the last LPS injection further amplified this reduction at day 22 (*3-way ANOVA on log transformed data, treatment effect, p=0.0108,* **Supplementary Table 7**).

In summary, we found that WT and KO animals exhibited similar pathological symptoms acutely after injections of Poly(I:C) and LPS. The alternating protocol of Poly(I:C) and LPS injection used in this study, while mitigating the development of tolerance, did not reveal a potentiation of the effects of successive injections. Furthermore, loss of Parkin function did not significantly modulate this response.

### Minimal peripheral inflammation persists at 6 months after a combined Poly(I:C)/LPS treatment

Within the framework of this model and given the potential implication of chronic inflammation in PD, we tested the hypothesis that the alternating Poly(I:C)/LPS protocol may lead to chronic peripheral inflammation, especially in Parkin KO mice. We maintained the mice for 6 months after the last injection **(Fig. 3A)** after which we quantified the levels of fecal calprotectin and lipocalin. We observed that fecal calprotectin levels were globally higher in Parkin KO mice compared to WT (*2-way ANOVA, genotype effect p=0.0378,* **Supplementary Table 8**), but that the treatment had no effect **(Fig. 3B)**. Notably, the detected calprotectin levels were globally low, and remained below those measured at the end of the acute phase (**Fig. 2D**). Similarly, fecal lipocalin-2 levels at 6 months were lower than during the acute phase and were no longer significantly elevated in the treated groups.

**Figure 3:**
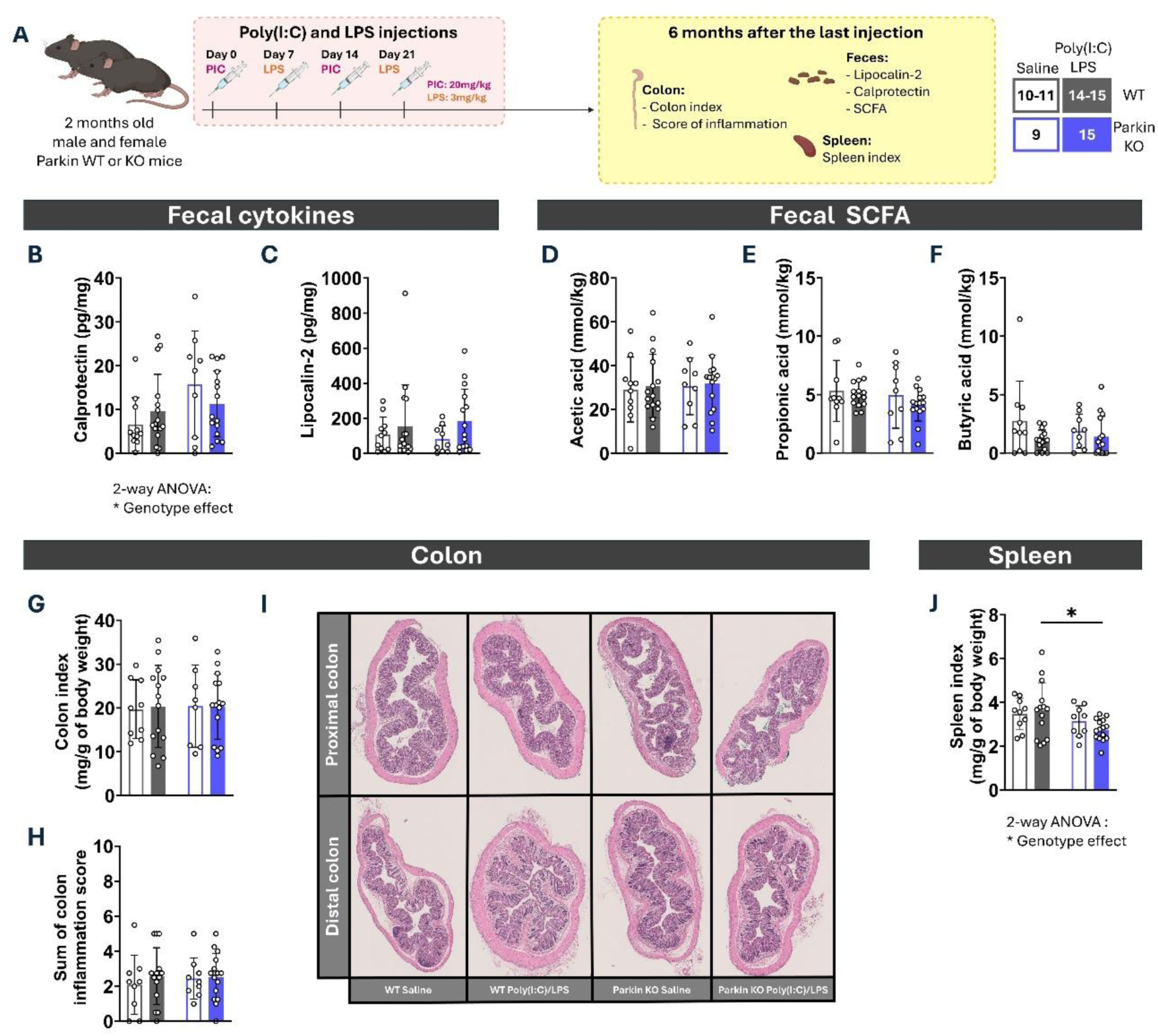
Absence of marked peripheral inflammation 6 months after the combined Poly(I:C)/LPS treatment. **(A)** Schematic illustration of the experimental design. WT and Parkin KO mice received alternate Poly(I:C)/LPS injections. The level of peripheral inflammation was evaluated 6 months after the last injection by measuring diverse inflammation markers in the colon, feces and spleen. **B-G,** fecal cytokines were detected by ELISA. **(B)** Calprotectin was detected at a higher overall level in the feces of Parkin KO mice while **(C)** Lipocalin-2 was not modulated by genotype or treatment. **D-F,** the quantity of fecal SCFA were measured by gas chromatography. **(D)** Acetic acid, **(E)** propionic acid and **(F)** butyric acid do not appear to be modulated by genotype or treatment. **(G)** The colon index, calculated as colon mass/mouse body mass, was similar between the groups. **(H)** Histopathological analysis of the colon does not show elevated inflammation scores. **(I)** Representative images for proximal and distal colon. **(J)** The spleen index, calculated as spleen mass/mouse body mass, is lower in Parkin KO mice and particularly when they received the Poly(I:C)/LPS treatment. For panels **B-H and J**, bars represent the mean ± SD of the group and dots represent individual animals. Grey and blue colors are used respectively for WT and Parkin KO mice. Bars corresponding to saline are unfilled and outlined, while bars corresponding to Poly(I:C)/LPS are filled. Statistical significance was assessed using a 2-way ANOVA (* p<0.05), the results of which are presented in **Supplementary Table 8.** It was followed by Šídák’s multiple comparisons test. Significant comparisons are indicated above the bars (* adjusted p<=0.05). Transformation of the data before testing was done when necessary. n=10-11 for WT Saline, n=14-15 for WT Poly(I:C)/LPS, n=9 for Parkin KO Saline and n=15 for Parkin KO Poly(I:C)/LPS, from 2 independent cohorts.

Given that fecal SCFA levels have been reported to be reduced in people suffering with PD (Aho et al., 2021; Chen et al., 2022; Nishiwaki et al., 2024; X. Yang et al., 2022), we also quantified fecal SCFA in the feces 6 months after the last injection. Acetic acid, propionic acid and butyric acid levels were comparable across groups **(Fig. 3D-F).** Consistent with these findings, there was no evidence of ongoing colonic inflammation 6 months after the last injection, indicated by an unchanged colon index **(Fig. 3G)** and an absence of inflammatory indicators revealed by a histopathological analysis of the proximal and distal colon **(Fig. 3H & I)**.

Systemic inflammation could also modulate cell number in the spleen, a secondary lymphoid organ. Here, we observed an overall decrease in spleen index in Parkin KO mice (*2-way ANOVA on log transformed data, genotype effect p=0.0323*, **Supplementary Table 8**) **(Fig. 3J)**, with a significant decrease in Parkin KO mice receiving Poly(I:C)/LPS compared to WT mice receiving the same treatment (*2-way ANOVA on log transformed data, followed by Šídák’s multiple comparisons test, adjusted p value=0.0365*). Although uncommon, splenic involution has been reported in the context of autoimmune diseases such as coeliac diseases (Bullen et al., 1981; Di Sabatino, 2013; Sabatino et al., 2006; Wardrop et al., 1975) and may reflect progressive impairment or exhaustion of splenic immune function associated with persistent systemic inflammation.

Elevated levels of pro-inflammatory cytokines and chemokines in the serum have been frequently reported in PD cohorts (Brodacki et al., 2008; Zimmermann and Brockmann, 2022). We therefore assessed a 32-plex panel of cytokines and chemokines in one cohort of mice 6 months after the treatment **(Suppl. Fig. 2A)**. Of the 32 analytes measured, the majority were unaffected by either treatment or genotype **(Suppl. Fig. 2B)**. The only significant differences were a decrease in the levels of IL-15 in Parkin KO mice, particularly marked in saline group (*2-way ANOVA on log transformed data, genotype effect p=0.0292,* **Supplementary Table 8,** *followed by Šídák’s multiple comparisons test, adjusted p value=0.0390*) **(Suppl. Fig. 2C),** as well as a genotype and treatment-dependent changes in the levels of LIX (CXCL5), as revealed by a significant interaction between genotype and treatment (*2-way ANOVA, interaction effect p=0.0347,* **Supplementary Table 8**) **(Suppl. Fig. 2D**).

Together these data indicate that 6 months after the alternate injection of Poly(I:C) and LPS, there is a minimal inflammation remaining in the periphery of both WT and Parkin KO mice.

### Motor functions and the morphological integrity of the dopamine system remain unaltered following the Poly(I:C)/LPS treatment

The characteristic motor symptoms of PD are linked to a loss of midbrain dopaminergic neurons. Since previous work using a chronic LPS injection model reported the onset of motor dysfunctions approximatively 4 months after the final injection(Liu et al., 2008), we hypothesized that alterations in the DA system and motor functions might similarly emerge in our alternating Poly(I:C)/LPS model 4 months after the last injection, with potentially larger changes in Parkin KO mice.

We first examined spontaneous locomotor activity using an open field test **(Fig. 4A)**. Total movement time **(Fig. 4B),** time spent in the center of the field **(Fig. 4C**), time spent in the periphery **(Fig. 4D)**, total distance traveled **(Fig. 4E)** and number of vertical episodes **(Fig. 4F)** were not significantly different across treatments and genotypes. We conclude that spontaneous locomotor activity was not impacted by the treatment.

**Figure 4:**
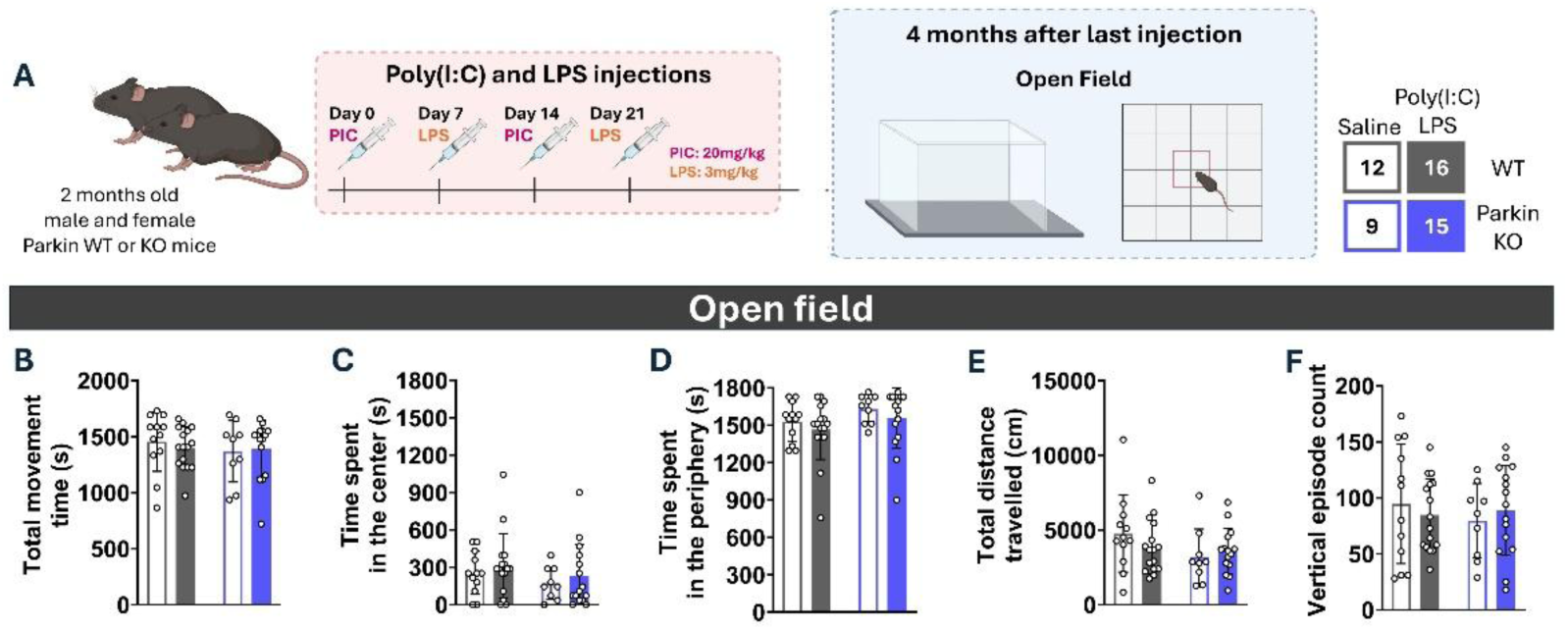
Repeated Poly(I:C)/LPS treatments do not change the exploratory locomotor activity of WT or Parkin KO mice. **(A)** Schematic illustration of the experimental design. WT and Parkin KO mice received alternate Poly(I:C)/LPS injections. Exploratory locomotion was assessed 4 months after the last injection. Mice were placed in an open field for 30 min. The time that the animal spent **(B)** in movement, **(C)** in the center **(D)** or in the periphery, **(E)** their total distance travelled and **(F)** the count of vertical episodes, were not affected by genotype or the Poly(I:C)/LPS treatment. For panels **B-F**, bars represent the mean ± SD of the group and dots represent individual animals. Grey and blue are used respectively for WT and Parkin KO mice. Bars corresponding to saline are unfilled and outlined, while bars corresponding to Poly(I:C)/LPS are filled. n=12 for WT Saline, n=16 for WT Poly(I:C)/LPS, n=9 for Parkin KO Saline and n=15 for Parkin KO Poly(I:C)/LPS, from 2 independent cohorts.

The mice were also evaluated in more demanding tasks assessing motor skills and coordination. **(Fig. 5A)**. In the pole test, no statistically significant difference was observed between genotypes or treatment for the time required for the animals to orient themselves facing in a downward direction **(Fig. 5B)** and for the time required to climb down the pole **(Fig. 5C)**. In the rotarod task, the animals first underwent a training period and the number of trials required to reach a stable performance at 4 rpm was recorded. The Poly(I:C)/LPS-treated mice required more trials to reach criterion (*2-way ANOVA, treatment effect p=0.0470,* **Supplementary Table 9**) **(Fig. 5D)**. However, the treated animals progressed similarly to the controls over the 9 test sessions (*3-way ANOVA on square root transformed data, time effect, p<0.0001,* **Supplementary Table 9**), with equivalent latencies to fall (**Fig. 5E)**.

**Figure 5:**
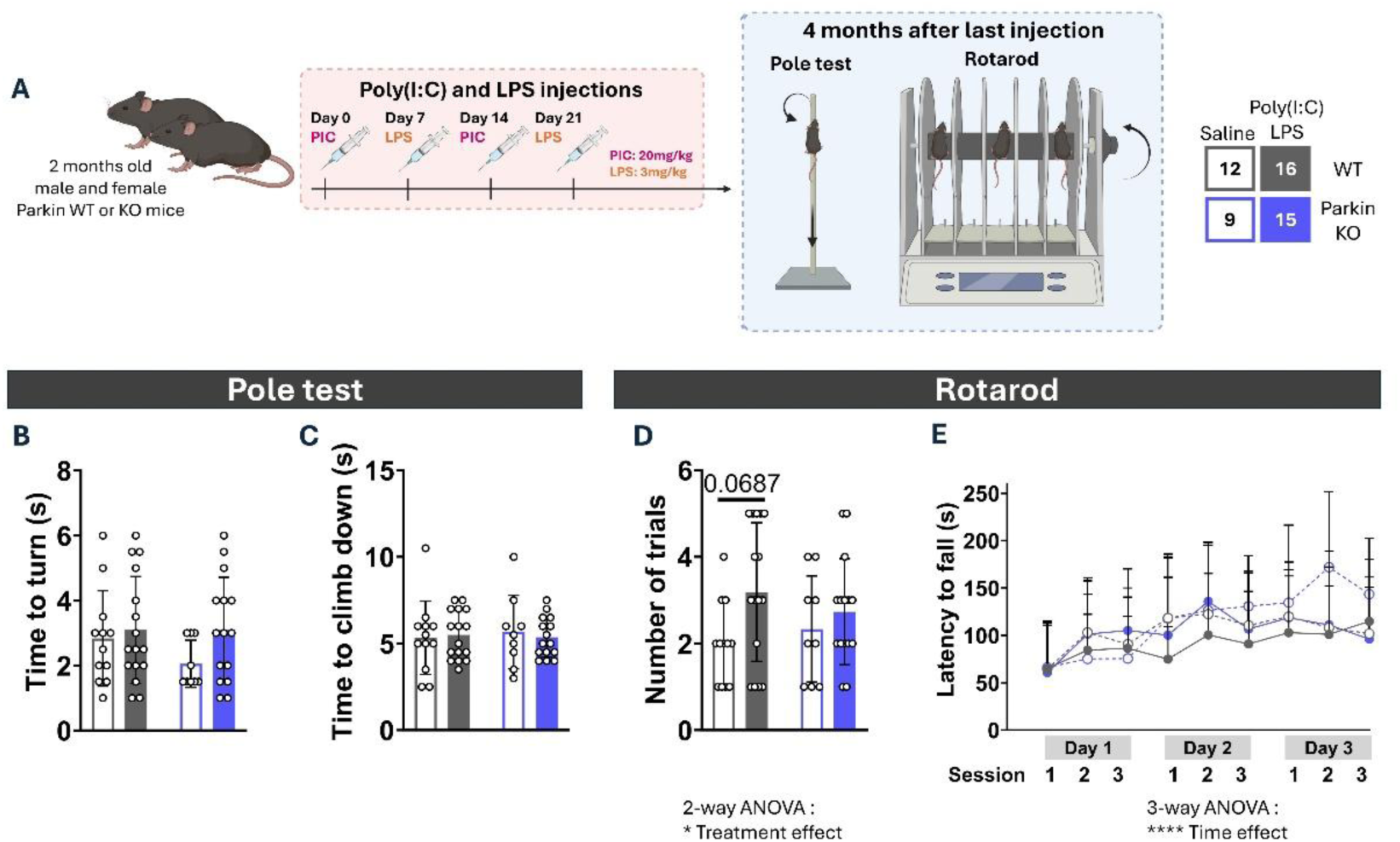
Repeated Poly(I:C)/LPS treatments do not change the motor capacities of WT or Parkin KO mice. **(A)** Schematic illustration of experimental design. WT and Parkin KO mice received alternate Poly(I:C)/LPS. Motor capacities were assessed 4 months after the last injection, using the pole test and the rotarod test. **B-C**, Mice were tested on a pole test for 3 trials. The first trial was considered as a habituation, and the results are presented as the mean of the last two trials. **(B)** The time required to turn and **(C)** the time necessary for the mouse to climb down completely are not impacted by the genotype or the treatment. **(D)** Mice were pre-trained on the rotarod at a constant speed of 4 rpm to reach a stable performance and the number of assays were noted. Mice that received Poly(I:C)/LPS injections need more trials, particularly WT mice. Statistical significance was assessed using a 2-way ANOVA (* p<=0.05), the results of which are presented in **Supplementary Table 9.** It was followed by Šídák’s multiple comparisons test and significant comparisons are indicated above the bars (adjusted p>0,05, nonsignificant). **(E)** The day after training, mice were tested for 3 consecutive days on the rotarod with 3 sessions per day, each session consisting of accelerated rotations of 4 to 40 revolutions per minute. The time spend by the mice on the rotarod before falling is represented. Each group showed learning and no differences between these groups were observed. Statistical significance was assessed using 3-way ANOVA (**** p<0.0001), the results of which are presented in **Supplementary Table 8**. For panels **B-D,** bars represent the mean ± SD of the group and dots represent individual animals. Grey and blue are used respectively for WT and Parkin KO mice. Bars corresponding to saline are unfilled and outlined, while bars corresponding to Poly(I:C)/LPS are filled. **E,** Dots with error bars depict the mean ± SD for each group. Dotted lines represent saline-treated controls, and solid lines represent treated animals. n=12 for WT Saline, n=16 for WT Poly(I:C)/LPS, n=9 for Parkin KO Saline and n=15 for Parkin KO Poly(I:C)/LPS, from 2 independent cohorts.

We next examined the morphological integrity of the dopaminergic system. Six months after the last injection, we quantified the density of TH-positive axonal processes in the dorsal and ventral sectors of the striatum. **(Fig. 6A & B)**. The surface area of TH immunoreactivity **(Fig. 6C & G)** and of DAT immunoreactivity **(Fig. 6D & H)** in the ventral and dorsal striatum was unchanged by treatment or genotype. The intensity of TH and DAT immunoreactivity was also not significantly different (**Fig. 6E-F & I-J)**. Together these findings indicate that the DA system did not undergo detectable degeneration, corroborating the absence of motor impairments in these animals.

**Figure 6:**
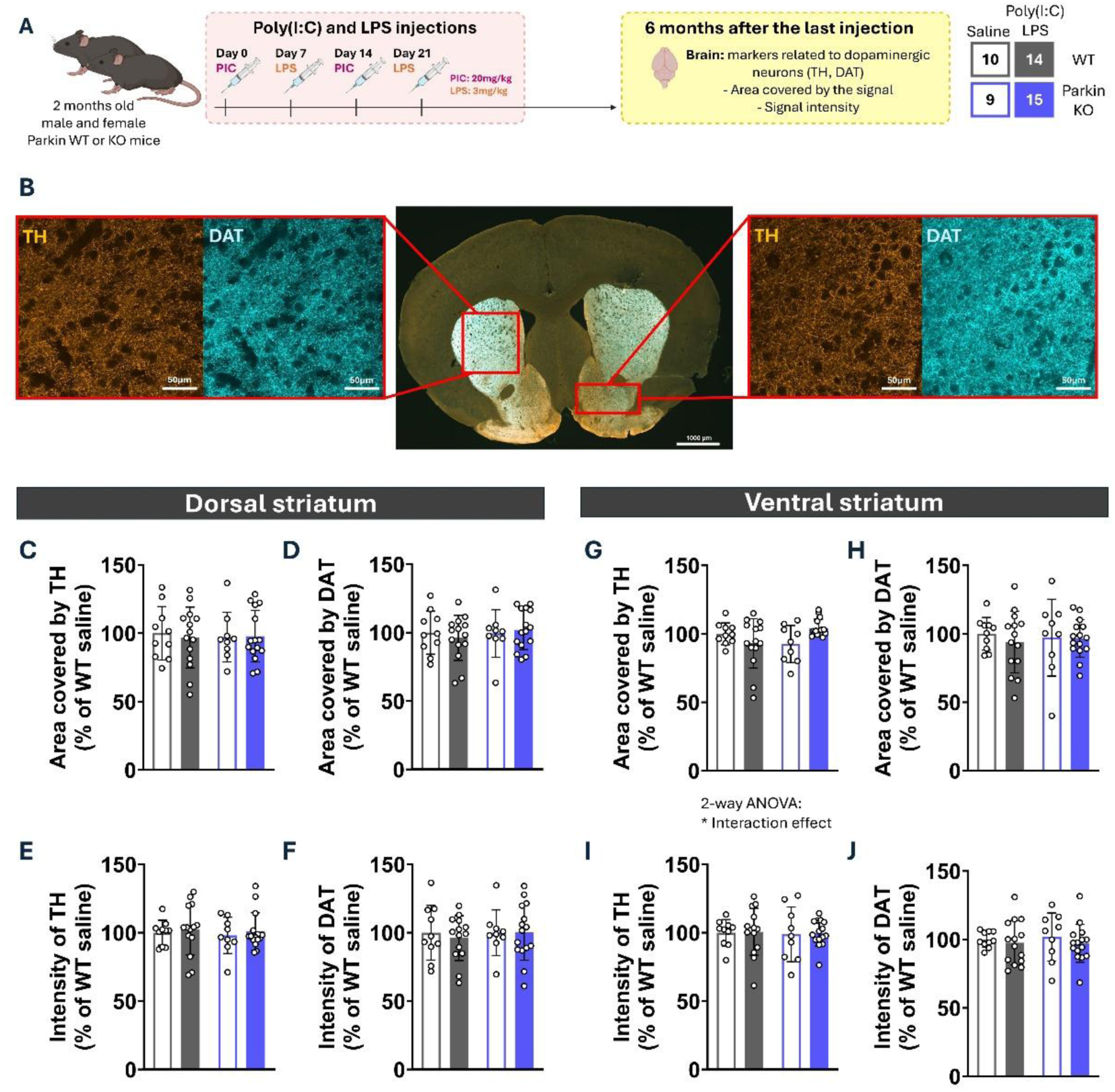
Repeated Poly(I:C)/LPS treatments do not impact dopaminergic neuron markers in the striatum. **(A)** Schematic illustration of the experimental design. WT and Parkin KO mice received alternate Poly(I:C)/LPS injections. The impact of this treatment on dopaminergic innervation in the striatum was evaluated 6 months after the last injection by measuring TH and DAT markers on 3 striatal slices. **(B)** Representative images of immunohistofluorescence in striatal brain slices showing expression of TH (orange) and DAT (cyan blue) in the whole slice (scale bar: 1000 µm) and dorsal or ventral striatum (scale bar: 50 µm). Images were acquired with a confocal microscope as a large image at X20. **C-F**, in the dorsal striatum, the area covered by **(C)** TH or **(D)** DAT signals and the intensity of **(E)** TH or **(F)** DAT signals are similar among the groups. **G-J,** In the ventral striatum, **(G)** only the area covered by TH was modulated by both the treatment and genotype, **(H, I &J)** with other parameters showing no difference between groups. For panels **C-I,** bars represent the mean ± SD of the group and dots represent individual animals. Grey and blue are used respectively for WT and Parkin KO mice. Bars corresponding to saline are unfilled and outlined, while bars corresponding to Poly(I:C)/LPS are filled. Statistical significance was assessed using a 2-way ANOVA on log transformed data (* p<=0.05), the results of which are presented in **Supplementary Table 10.** n=10 for WT Saline, n=14 for WT Poly(I:C)/LPS, n=9 for Parkin KO Saline and n=15 for Parkin KO Poly(I:C)/LPS, from 2 independent cohorts.

Although PD is accompanied by the degeneration of dopaminergic neurons in the substantia nigra, it also affects other parts of the central and peripheral nervous system, including the autonomic nervous system (Borghammer and Van Den Berge, 2019; Skjærbæk et al., 2026).

For example noradrenergic neurons in the stellate ganglia can be affected (Orimo et al., 2005). Here we examined sympathetic (noradrenergic) and parasympathetic (cholinergic) markers in the stellate ganglion of Poly(I:C)/LPS-treated mice **(Suppl. Fig. 3A).** TH immunoreactivity, used to label noradrenergic fibers, was not significantly affected 6 months after the last injection of Poly(I:C)/LPS. **(Suppl. Fig. 3B-D)**. However, the levels of choline acetyltransferase (ChAT) were globally lower after Poly(I:C)/LPS injections (*2-way ANOVA, treatment effect p=0.0217,* **Supplementary Table 10**). A genotype effect was also detected, with reduced ChAT immunoreactivity in Parkin KO mice (*2-way ANOVA, genotype effect, p=0.0065*) **(Suppl. Fig. 3E)**. We also found a significant genotype effect in Poly(I:C)/LPS treated mice, revealing a larger decrease in ChAT immunoreactivity in the Parkin KO tissue (*2-way ANOVA followed by Šídák’s multiple comparisons test, WT Poly(I:C)/LPS vs Parkin KO Poly(I:C)/LPS p=0.0449*). However, the area covered by the ChAT immunoreactive signal was unchanged **(Suppl. Fig. 3F)**, suggesting reduced ChAT levels, but not necessarily neuronal loss.

Together, these data suggest that although Poly(I:C)/LPS treatment does not detectably compromise the integrity of DA system, as supported by the absence of motor symptoms at this stage, it may nonetheless impact the peripheral nervous system, particularly in Parkin KO mice.

### Repeated Poly(I:C)/LPS exposure induce minor changes in glial cells

The lack of alteration of DA system in these mice can be hypothesized to be due to an insufficient level of chronic brain inflammation. We tested his possibility by evaluating microglial and astrocytic markers, often reported to be elevated in the postmortem PD brain (Gerhard et al., 2006). To this end, we performed immunohistochemical labelling against the microglial marker Iba1, on striatal **(Fig. 7A)** and mesencephalic **(Fig. 7B)** brain sections examined in 3D confocal stacks. No significant difference was observed in the number of microglia present in the dorsal striatum **(Fig. 7C)**, ventral striatum **(Fig. 7F)**, SN **(Fig. 7I)** and VTA **(Fig. 7L).** Furthermore, the intensity of Iba1, known to increase during microglial activation, was only modestly increased by the Poly(I:C)/LPS treatment in the dorsal striatum (*2-way ANOVA, treatment effect, p=0.0266,* **Supplementary Table 11**), mainly in the WT mice (*2-way ANOVA, WT Saline vs WT Poly(I:C)/LPS, p=0.0295,* **Supplementary Table 11**) **(Fig. 7D)**. The other regions do not show significant modulation of Iba1 intensity **(Fig. 7G, J & M)**.

**Figure 7:**
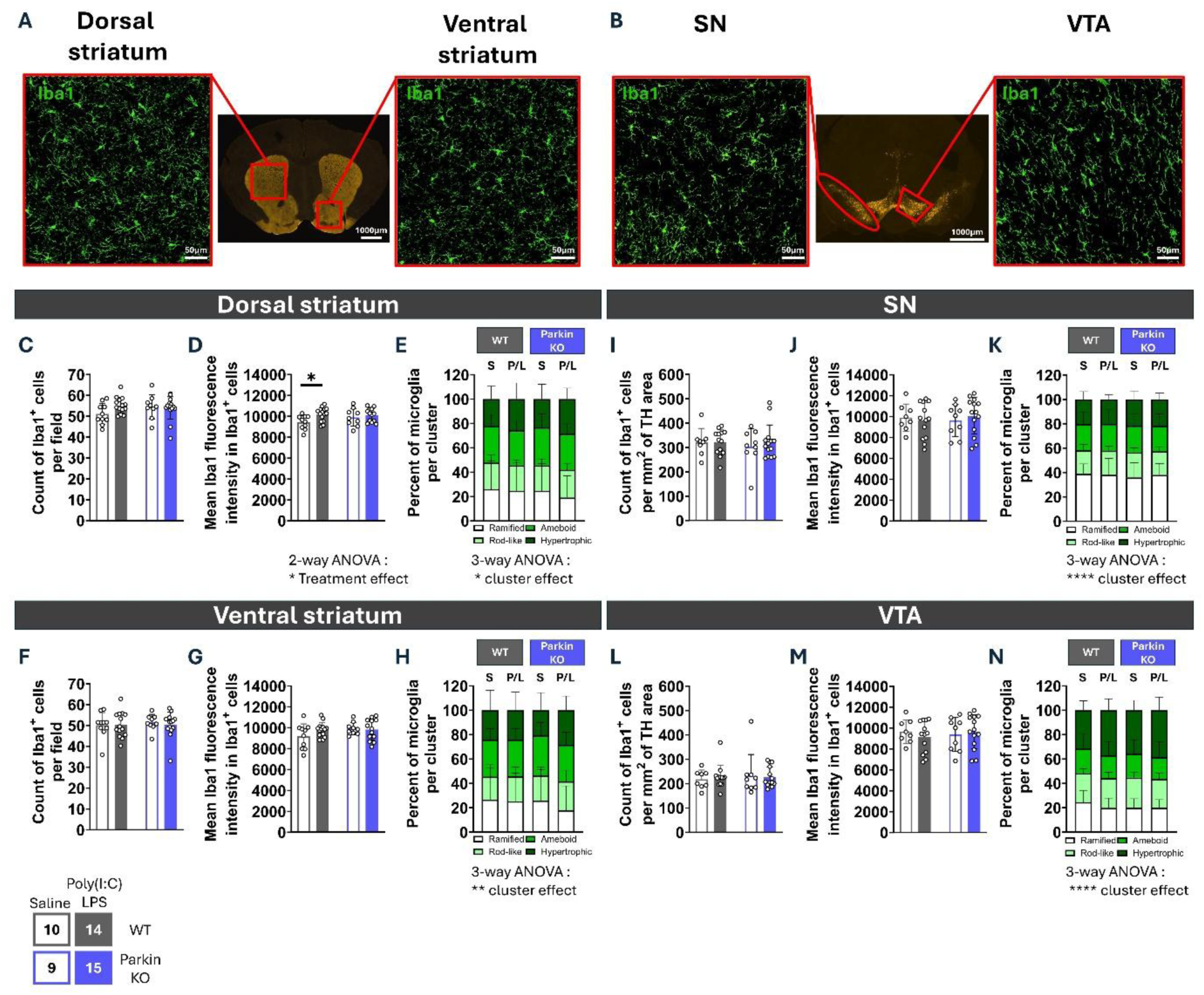
Repeated injections of Poly(I:C)/LPS do not change microglial density but induce minor changes in microglial Iba1 intensity in the dorsal striatum. A-B, Representative immunohistofluorescence images acquired with a confocal microscope and showing expression of TH (orange) in a whole brain slice (large image at X20, scale bar: 1000 µm) and expression of Iba1 (green) in 3D reconstructed images for the respective regions (X40, scale bar: 50 µm). **(A)** Images acquired in the striatum (dorsal and ventral). **(B)** Images acquired in the mesencephalon (SN and VTA). **C-E**, In the dorsal striatum, **(C)** the number of Iba1^+^ cells counted in the field was similar among the groups. **(D)** The mean fluorescence intensity of Iba1 measured in the Iba1^+^ cells was increased by the Poly(I:C)/LPS treatment while **(E)** the morphology of the microglia was unchanged. **F-H,** the three parameters were not modulated by the genotype or treatment in the ventral striatum, **I-K** or in the TH-positive areas in the SN and **L-N** VTA. For panels **C-D, F-G, I-J, L-M,** bars represent the mean ± SD of the group and dots represent individual animals. Grey and blue are used respectively for WT and Parkin KO mice. Bars corresponding to saline are unfilled and outlined, while bars corresponding to Poly(I:C)/LPS are filled. Statistical significance was assessed using 2-way ANOVA (* p<=0.05), the results of which are presented in **Supplementary Table 11.** It was followed by Šídák’s multiple comparisons test and significant comparisons are indicated above the bars (* adjusted p<=0.05). Transformation of the data before testing has been done when necessary. For panels **E, H, K, N,** stacked bars show the mean frequency ± SD of four morphology types inside a group. Morphology clusters are distinguished by shades of green: white for ramified, light green for rod-like, medium green for ameboid and dark green for hypertrophic. S: Saline, P/L: Poly(I:C)/LPS. Statistical significance was assessed using 3-way ANOVA (* p<=0.05, ** p<0.01, **** p<0.0001), the results of which are presented in **Supplementary Table 11.** Transformation of the data before testing was done when necessary. Images not meeting the predefined quality criteria were excluded from the analysis. n=8-10 for WT Saline, n=13-14 for WT Poly(I:C)/LPS, n=9 for Parkin KO Saline and n=13-15 for Parkin KO Poly(I:C)/LPS, from 2 independent cohorts.

We further examined microglial morphology. Microglia were grouped into four morphological clusters **(Fig. 7E, H, K, N)** corresponding to activation states: ramified (homeostatic), rod-like (intermediate/reactive), amoeboid (highly activated/phagocytic), and hypertrophic (reactive/inflammatory) phenotypes (Green and Rowe, 2024; Kim et al., 2024a; Reddaway et al., 2023). A 3-way ANOVA showed a significant effect of morphological clusters (*3-way ANOVA after log or square transformation, cluster effect p=0.0466 in dorsal striatum, p=0.0086 in ventral striatum, p<0.0001 in SN, p=<0.0001 in VTA*). Post hoc comparisons confirmed significant differences between clusters, whereas no differences were observed between genotypes or treatment conditions.

Healthy astrocytes offer neuroprotection to neurons (Chiareli et al., 2021), but in response to stress or injury, they become reactive, often linked to an elevation of GFAP expression. We investigated in the striatum **(Fig. 8A)** and mesencephalon **(Fig. 8B)** the number of GFAP^+^ cells **(Fig. 8C, F, I & L)**, the volume of GFAP-immunoreactive signal **(Fig. 8D, G, J & M)** and the intensity of this GFAP immunoreactivity **(Fig. 8E, H, K & N)**. Overall, the only significant changes were observed in the SN and VTA. In the SN, we detected a decrease in the number of GFAP-positive cells in brain slices from Poly(I:C)/LPS-treated mice **(Fig. 8I)**. This decrease was mainly observed in WT mice, as confirmed by a significant interaction between treatment and genotype in the 2-way ANOVA. There was also a tendency for a reduced number of GFAP-positive cells in non-treated Parkin KO mice (*2-way ANOVA, Treatment effect p=0.0341, Interaction effect p=0.0641, followed by Šídák’s multiple comparisons test, WT Saline vs WT Poly(I:C)/LPS adjusted p=0.0145*, **Supplementary Table 12**). The volume of GFAP signal was also reduced in the SN of WT mice after exposure to Poly(I:C)/LPS (*2-way ANOVA, Interaction effect p=0.0117, followed by Šídák’s multiple comparisons test, WT Saline vs WT Poly(I:C)/LPS adjusted p=0.0150, WT Saline vs KO Saline adjusted p=0.0318*, **Supplementary Table 12**) **(Fig. 8J)**. In the VTA, the number of GFAP astrocytes was also reduced, by Poly(I:C)/LPS both, particularly in WT animals (*2-way ANOVA, Treatment effect p=0.0584, Interaction effect interaction effect p=0.0273, followed by Šídák’s multiple comparisons test, WT Saline vs WT Poly(I:C)/LPS adjusted p=0,0113,* **Supplementary Table 12**) **(Fig. 8L)**. Moreover, GFAP immunoreactivity in the VTA was modestly decreased by the Poly(I:C)/LPS treatment (*2-way ANOVA, treatment effect, p=0.0228,* **Supplementary Table 12**) **(Fig. 8N)**.

**Figure 8:**
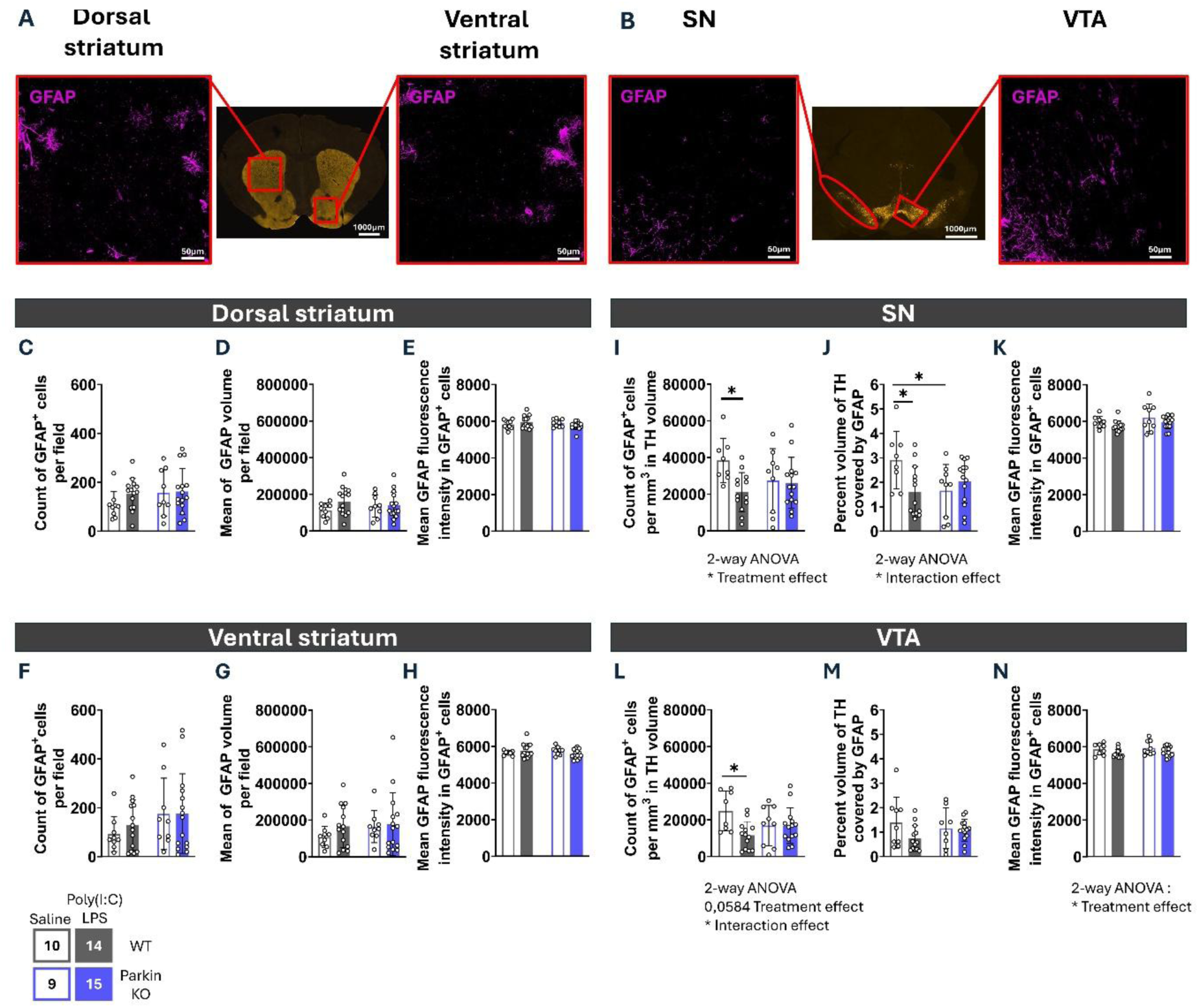
Repeated injections of Poly(I:C)/LPS induce a modest decrease of the number of GFAP positive cells in the SN and of the intensity of GFAP in VTA astrocytes. **A-B,** Representative immunohistofluorescence images acquired with a confocal microscope and showing expression of TH (orange) in a whole brain slice (large image at X20, scale bar: 1000 µm) and expression of GFAP (magenta) in 3D reconstructed images for the respective regions (X40, scale bar: 50 µm). **(A)** Images acquired in the striatum (dorsal and ventral). **(B)** Images acquired in the mesencephalon (SN and VTA). **C-E,** In the dorsal striatum, **(C)** the number of astrocytes, counted as positive for DAPI and within the GFAP-positive area, **(D)** the volume occupied by the GFAP signal in the field and **(E)** the mean fluorescence intensity of GFAP measured in the GFAP^+^ volume was not modulated by the treatment or the genotype. **F-H,** same for the ventral striatum. **I-K,** In the SN, **(I)** the number of astrocytes was higher in the WT saline group compared to the others, while **(J)** the volume and **(K)** intensity of GFAP remained the same among the other groups. **L-N,** In the VTA, **(L)** the number of GFAP-positive cells and **(M)** the volume of GFAP-positive signal was similar among the groups. However, **(N)** there was a small increase of GFAP immunoreactivity in mice treated with Poly(I:C)/LPS. **C-N,** bars represent the mean ± SD of the group and dots represent individual animals. Grey and blue are used respectively for WT and Parkin KO mice. Bars corresponding to saline are unfilled and outlined, while bars corresponding to Poly(I:C)/LPS are filled. Statistical significance was assessed using a 2-way ANOVA (* p<=0.05), the results of which are presented in **Supplementary Table 12.** It was followed by Šídák’s multiple comparisons test and significant comparisons are indicated above the bars followed (* adjusted p<=0.05). Images not meeting the predefined quality criteria were excluded from the analysis. n=8-10 for WT Saline, n=13-14 for WT Poly(I:C)/LPS, n=9 for Parkin KO Saline and n=14-15 for Parkin KO Poly(I:C)/LPS, from 2 independent cohorts.

These data indicate that the repeated Poly(I:C)/LPS treatment used in the present study induced only a modest overall long-term impact on glial cells. This effect was primarily characterized by a small increase in microglial cell activation at 6 months post-insult, most notably in the dorsal striatum. This was accompanied by reduced density of GFAP-positive astrocytes in WT mice, with similar reductions in both challenged and unchallenged KO mice in the SN. The subtle changes in the brain observed six months after the final injection warrant longer-term follow-up in future studies to determine whether they progress to sustained neuroinflammation or dopaminergic dysfunction in this model.

## Discussion

PD is a multifactorial disorder involving perturbations of both the central and peripheral nervous systems, and the mechanisms driving its onset are still not fully elucidated. Inflammation is thought to play a central role in its pathophysiology, and accumulating evidence suggests that infections may act as a potential trigger of the disease. Studying these mechanisms requires animal models in which the contribution of acute and chronic inflammation can be manipulated, yet current inflammatory models are typically limited by their highly acute nature or because they use complex and time-consuming experimental procedures. Here we aimed to develop a more accessible model that recapitulates both central and peripheral inflammatory features without requiring a multi-month protocol.

Repeated systemic LPS administration is known to induce tolerance (Kehl et al., 2004; Liu et al., 2008; Musaelyan et al., 2018; Y. Yang et al., 2022). In line with previous studies, initial experiments with repeated LPS confirmed the development of a gradual habituation, even in Parkin KO mice, despite their heightened sensitivity to inflammatory stress (Matheoud et al., 2016; Mouton-Liger et al., 2018). Strategies using low-dose LPS reduce tolerance and potentiate dopaminergic neuron loss (Liu et al., 2008) but fail to establish chronic inflammation or robust motor symptoms, even in Parkin KO animals (Frank-Cannon et al., 2008). In PD, both bacterial and viral infections are considered as potential triggers (Smeyne et al., 2020). Poly(I:C) mimics viral infection while inducing limited tolerance upon successive injections (Cunningham et al., 2007). Based on these observations, we evaluated a novel alternating injection protocol of Poly(I:C) and LPS, thereby reducing tolerance and potentially maintaining peripheral inflammatory responses over an extended period. To our knowledge, this is the first study evaluating a potential disease model based on the paradigm of alternating systemic administration of dual PAMPs.

Using this paradigm, a consistent body weight loss was observed after each injection, in contrast with protocols using a single PAMP. Given the systemic nature of this protocol, we examined whether repeated Poly(I:C)/LPS administration induced alterations in peripheral organs, with particular emphasis on the gastrointestinal tract, which is commonly affected in PD. Fecal lipocalin-2, elevated in both acute and chronic intestinal dysbiosis (Thorsvik et al., 2017; Yadav et al., 2022; Zollner et al., 2021), is not consistently found elevated in the feces of people suffering from PD. However, its modulation has been correlated with gut permeability changes and alterations in microbiota composition (Aho et al., 2021), relevant to PD. We found marked increase of lipocalin-2 acutely after PAMP injections, to a similar level in Parkin KO compared to WT mice. At later stages, baseline lipocalin-2 levels declined, which may reflect an age-related trend observed in both human and rodent studies (Qiu et al., 2021; Tarassishin et al., 2025). Although the overall comparison was not statistically significant, it is important to note that most animals maintained high fecal lipocalin levels in the Poly(I:C)/LPS treated group, and particularly in Parkin KO mice. The apparent presence of subgroups within the dataset, together with the limited sample size, prevents determining whether this trend reflects a true treatment effect. This pattern is consistent with a potential influence of a reduction of microbiome diversity or an impaired intestinal barrier integrity induced by the inflammatory insult and the genotype (Aho et al., 2021), with some individuals being more susceptible than others. Interestingly, two studies have suggested that Parkin deficiency may contribute to disruption of gut homeostasis (Cao et al., 2020; Feltzin et al., 2019). However, we did not observe any on-going colon damage or inflammation by immunohistopathology at 6 months post challenge. However, it should be noted that immune-cell infiltration may decline below detection threshold after cessation of the inflammatory insult. Moreover, residual or low-grade intestinal inflammation may have been missed. Furthermore, a reduced inflammatory cell infiltration in the gut does not necessarily imply that the tissue has returned to its baseline state. Chronic inflammation commonly fluctuates over time, whereas its structural consequences, such as extracellular-matrix deposition and fibrotic remodeling may persist or progress. Additional assessment of tissue-remodeling markers is needed to distinguish fully resolved inflammation from persistent structural changes in the absence of active inflammation. Moreover, people with PD also exhibit elevated fecal calprotectin levels (Aho et al., 2021; Dumitrescu et al., 2021; Hor et al., 2022) and reduced concentrations of fecal SCFA (Aho et al., 2021; Chen et al., 2022; Nishiwaki et al., 2024; X. Yang et al., 2022), both of which have been associated with disease progression. In contrast to lipocalin-2, we found that calprotectin levels remained stable immediately after injections, suggesting that the treatment did not induce detectable neutrophil-associated intestinal response (Prata et al., 2016; Thorsvik et al., 2017). Given the i.p. route of administration, this finding is not unexpected, as the intestinal lumen and mucosa were not directly exposed to the inflammatory stimuli. Nevertheless, the absence of an early increase in fecal calprotectin does not exclude the induction of systemic inflammation or more subtle intestinal effects. At the long-term time point, fecal calprotectin levels were lower overall, although Parkin KO mice retained slightly higher levels than WT mice, irrespective of the inflammatory challenge. Similar decreases have been reported in children with high fecal *E. coli* abundance (Orivuori et al., 2015), raising again the possibility of microbiota-driven modulation. However, long-term levels of SCFA were similar between the groups. Taken together, our results indicate that alternating dual PAMP exposure may promote a chronic low-grade inflammation following the initial challenge, often associated with neurodegenerative disease, rather than a dense neutrophil infiltration typically associated with acute intestinal inflammation. Future studies incorporating longitudinal microbiota profiling, intestinal barrier function assays and additional inflammatory markers will be important for determining whether the treatment induces persistent low-grade inflammation or longer-lasting tissue alterations. Evidence for such effects would be strengthened by convergent findings, including sustained cytokine changes, altered intestinal architecture or permeability, shifts in immune-cell populations, and increased tissue-remodelling markers such as collagen deposition or α-smooth muscle actin.

Despite these peripheral signatures, motor impairments did not emerge within four months after the last PAMP injection. However, conventional behavioral assays such as the open field test and the pole test may lack sensitivity to detect subtle deficits. It should be noted that measurable impairments in such tasks typically requires a substantial loss of the nigrostriatal DA in rodents. Interestingly, Poly(I:C)/LPS-treated animals displayed impaired learning on the rotarod. This suggests that neuroinflammation induced by alternating PAMPs may disrupt motor learning before affecting motor functions *per se*. This interpretation is consistent with previous reports showing that systemic administration of LPS or Poly(I:C) can impair cognitive functions and hippocampal activity (Garré et al., 2017; Morimoto et al., 2023; Schirmbeck et al., 2023). However, because rotarod performance is influenced by both motor coordination and procedural learning, additional cognitive and motor assays will be necessary to distinguish these effects.

In PD, early cognitive dysfunction is associated with more severe and faster progressing motor and non-motor symptoms including constipation and gastric dumping. Moreover, recent evidence suggests that oral and gut microbial profiles are associated with cognitive decline and disease progression in PD (Clasen et al., 2025). Thus, the chronic, low-grade intestinal effects associated with repeated dual PAMP exposure may contribute to cognitive dysfunction in this model, thereby mimicking PD-associated dementia rather than acute inflammatory insult. This increases the translational value of the model, as sustained, low-level systemic inflammatory signaling is hypothesized to progressively compromise brain function during the extended prodromal phase, as observed in human PD. Assessing cytokine expression, blood–brain barrier integrity, and microglial activation in brain regions involved in motor learning and cognition would help determine whether neuroinflammatory mechanisms contribute to the observed behavioral phenotype.

We did not detect significant changes in dopaminergic fibers in the striatum, at least based on TH immunoreactivity. Although we cannot exclude that subtle degeneration occurred in the substantia nigra, as observed by Franck-Cannon et al in their LPS repeated injection model (Frank-Cannon et al., 2008). Additional experiments including stereological counting of DA neuron cell bodies would be required to validate this. Although the central DA system did not appear to be affected in our model at this stage, the reduced expression of ChAT in the stellate ganglia points to a potential early impairment of cholinergic function in the periphery. This finding is of particular interest in light of evidence that LPS-challenged VAChT-knockdown mice exhibit increased susceptibility to inflammation and mortality, accompanied by elevated TNF-α, IL-1β, and IL-6 levels in the spleen and brain (Leite et al., 2016), suggesting that LPS exposure may disrupt cholinergic anti-inflammatory signaling. These observations are noteworthy given that PD is increasingly recognized as a multisystem disorder involving both central and peripheral cholinergic dysfunction (Bohnen et al., 2022; Pasquini et al., 2021). In addition to well-characterized central cholinergic deficits, enteric cholinergic abnormalities have been reported in patients with PD (Fedorova et al., 2017; Gjerløff et al., 2015), raising the possibility that early peripheral cholinergic alterations may emerge before, or contribute to, central neurodegenerative changes. Further investigation of the cholinergic system will be required to determine whether the reduction in ChAT expression reflects a broader impairment of cholinergic signaling. In particular, future studies should assess whether cholinergic alterations extend beyond the stellate ganglia to other peripheral tissues, including the enteric nervous system, as reported in patients with PD, as well as evaluate complementary markers of cholinergic neurons and function. Such analyses would help establish whether peripheral cholinergic dysfunction represents an early consequence of repeated inflammatory challenge or a mechanism linking systemic inflammation to PD-related pathogenesis

We did not observe a major change in microglial morphology or Iba1 expression 6 months after the end of the Poly(I:C)/LPS treatment. Only a modest increase of the intensity of Iba1 was observed in the dorsal striatum. Repeated administrations of LPS for a minimum five months were shown to induce an elevation of CD45 in the midbrain of WT and Parkin KO mice (Frank-Cannon et al., 2008; Liu et al., 2008). This suggests that a high number of inflammatory challenges may be necessary for a sustained change in microglial activity. Intriguingly, in another study using Parkin-deficient mice, a single LPS injection was shown to elicit a stronger microglial activation in Parkin KO compared to WT animals (Yan et al., 2023). In that study, the injections were performed at an older age, suggesting the possibility of age-dependent modulation of the microglial response in Parkin deficient-mice. We cannot exclude that repeated PAMP exposure in the present study lead to long term microglial tolerance (Kim et al., 2024b). But we must also note that Iba1 expression and the morphology of microglia do not represent the only indicators of microglia activation. In fact, previous work has shown that days following repeated LPS injections, microglia may return to baseline level of Iba1expression, while continuing to release pro-inflammatory mediators, including IL-1β (Bodea et al., 2014). Thus, the absence of sustained changes in Iba1 or microglial morphology would not exclude persistent microglial activation or altered function in the present model. In the context of neurodegenerative diseases, chronic low-grade inflammation may involve more modest or transient changes in Iba1 and cellular morphology while maintaining sustained alterations in cytokine release, immune signaling, or glial functions, compared to acute inflammatory context. Therefore, the findings in this model may reflect persistent functional dysregulation rather than overt, morphologically defined microgliosis.

Interestingly, we detected a significant decrease in GFAP-positive astrocytes in the SN and VTA, influenced both by treatment and Parkin deficiency. Astrocytes are central to neuronal survival (Chiareli et al., 2021) and are increasingly recognized as key contributors to neuronal vulnerability in PD (Booth et al., 2017; Miyazaki and Asanuma, 2020). Although reactive astrogliosis with elevated GFAP is frequently reported in PD models, our findings suggest a reduced number of astrocytes and/or of their activity. However, a decrease in GFAP-positive cells does not necessarily indicate astrocyte loss. It may instead reflect altered GFAP expression, changes in astrocytic reactivity, or a shift toward a less readily detectable cellular state. Interestingly, a decrease in GFAP expression has been reported in other models of neurodegenerative diseases including amyotrophic lateral sclerosis and Alzheimer’s disease, both linked to chronic neuroinflammation (Díaz-Amarilla et al., 2011; Kulijewicz-Nawrot et al., 2012). We can hypothesize that chronic inflammation may lead to a long-term functional impairment of astrocytes, associated with a phenotypic shift rather than a simple reactive state. A reduction of GFAP may also reflect impaired astrocyte proliferation, metabolic support, and antioxidant defense, consistent with evidence of mitochondrial dysfunction in Parkin-deficient astrocytes (Solano et al., 2008). Our team and others previously reported that glial cultures from Parkin KO mice harbor reduced number of astrocytes (Giguère et al., 2019; Solano et al., 2008). In an experimental autoimmune encephalomyelitis model, the number of GFAP-positive astrocytes was found significantly decreased in the white and grey matter of Parkin-deficient mice during the recovery phase (Cossu et al., 2021). Interestingly, we observed a decrease in IL-15 levels in Parkin KO mice in the present study. This finding may be of particular relevance because IL-15 plays dual roles: while required for CD8^+^ memory T cells and NK cell development (Goleij et al., 2025), it also promotes neuronal and astrocytic survival under stress by supporting mitochondrial functions (Lee et al., 2017; Li et al., 2018). Although the literature remains inconsistent regarding IL-15 levels in people with PD (Gangemi et al., 2003; Rentzos et al., 2007), the reduction observed in Parkin KO mice raises the possibility that impaired IL-15 signaling may contribute to astrocyte dysfunction. Otherwise, a similar decrease of GFAP has also been reported in organoids derived from patients carrying PRKN gene mutations (Kano et al., 2020). Moreover, reduced GFAP levels were observed in postmortem brains from PRKN mutation carriers (Kano et al., 2020), with an unchanged number of ALDH1L1-positive astrocytes, here again suggesting that astrocytic dysfunction rather than impaired proliferation predominates in absence of Parkin. Reduced GFAP levels could also be associated with an impaired capacity of astrocytes to regulate glutamate homeostasis and the blood-brain barrier (Kálmán, 2025). Assays examining glutamate uptake, lactate transport, and neurotrophic support would clarify whether astrocyte dysfunctions exist in the model developed in the present study.

Sex differences in PD have been largely reported (Cattaneo and Pagonabarraga, 2025), including potential differences in disease progression and clinical outcomes. In mice, females are less susceptible than males to DA neuron loss following LPS injection, requiring higher doses or repeated injections to achieve comparable neurodegeneration (Liu et al., 2008). However, one study showed that female Parkin mutant mice start to lose their DA neurons six months before their male counterparts (Rodríguez-Navarro et al., 2008), possibly due to a modulation of the cellular estrogen response in absence of Parkin. In the present study, no obvious sex-specific pattern emerged and results from both sexes were thus pooled. However, it must be noted that the small sample size prevented formal subgroup analyses and limited the statistical power to reliably evaluate sex-dependent effects.

Overall, the alternating Poly(I:C)/LPS injection paradigm established in the present study provides a useful proof-of-concept model for investigating the systemic and central consequences of repeated dual PAMP challenges. The protocol reliably induced early and robust body weight loss, accompanied with evidence of modest gut inflammation, particularly in the acute phase. In the brain, alternating Poly(I:C)/LPS exposure produced a measurable increase in microglial activation markers in the dorsal striatum, despite the absence of detectable morphological changes, and was associated with reduced GFAP expression in the SN, an effect that was more pronounced in the context of Parkin deficiency.

These findings highlight the sensitivity of the paradigm for detecting subtle, region-specific, and genotype dependent neuroimmune alterations that may precede overt neurodegeneration. Nevertheless, the absence of strong motor deficits or overt dopaminergic neurodegeneration indicates that, this model remains limited in recapitulating the central hallmarks of PD at 6 months post-challenge. However, this finding may also reflect the protracted nature of PD pathogenesis. As in human disease, in which peripheral and non-motor abnormalities can precede clinically evident dopaminergic dysfunction by many years, mice may require substantially longer follow-up after repeated inflammatory challenges before dopaminergic impairment becomes detectable. Thus, the current model may capture an early or prodromal stage rather than fully established PD pathology. Longitudinal studies incorporating extended behavioral, neurochemical, histological, and inflammatory assessments will be necessary to determine whether the observed peripheral and neuroimmune changes progress to dopaminergic dysfunction over time. Accordingly, the paradigm should not be regarded as a robust standalone model of PD, but rather as a tractable proof-of-concept platform for examining how repeated peripheral inflammatory stimuli interact with Parkin deficiency to shape early or non motor PD-like phenotypes. Applying this protocol in more penetrant Parkin variants as PrknR275W mice (Regoni et al., 2024), or in aged animals (Rocha et al., 2023) may accelerate or amplify central pathological outcomes and further improve the model’s translational relevance. Future longitudinal studies in these contexts may help establish whether chronic inflammatory exposure contributes to the gradual emergence of dopaminergic dysfunction, as observed in human PD.

## CRediT authorship contribution statement

**Amandine Even:** Writing – original draft, Writing – review & editing, Visualization, Validation, Methodology, Investigation, Software, Data curation, Formal analysis, Conceptualization. **Sriparna Mukherjee:** Writing – original draft, Writing – review & editing, Visualization, Validation, Methodology, Investigation, Conceptualization. **Nicolas Giguère:** Investigation, Software. **Morgane Brouillard-Galipeau:** Investigation. **Nathalia Luisa Oliveira:** Investigation. **Priyabrata Halder:** Investigation. **Claudie Beaulieu:** Investigation. **Romain Cayrol:** Investigation. **Nathalie Van Den Berge:** Investigation, Writing – review & editing. **Samantha Gruenheid:** Writing – review & editing, Resources, Conceptualization. **Louis-Eric Trudeau:** Writing – original draft, Writing – review & editing, Visualization, Validation, Supervision, Resources, Project administration, Methodology, Funding acquisition, Conceptualization.

## Acknowledgements

The authors thank Marie-Josée Bourque for her critical help with mouse breeding and Elie Bres for his help during sample collection. We would also like to thank Drs. Michel Desjardins, Heidi McBride, Jo Anne Stratton and Janelle Drouin-Ouellet for their critical feedback during the development of this project. The help of Lilia Rodriguez for the open science data sharing is also acknowledged.

## Data availability

The datasets, software/code, protocols, and lab materials used and/or generated in this study are listed in a Key Resource Table alongside their persistent identifiers at https://doi.org/10.5281/zenodo.20386044.

---

**Supplementary Figure 1:**
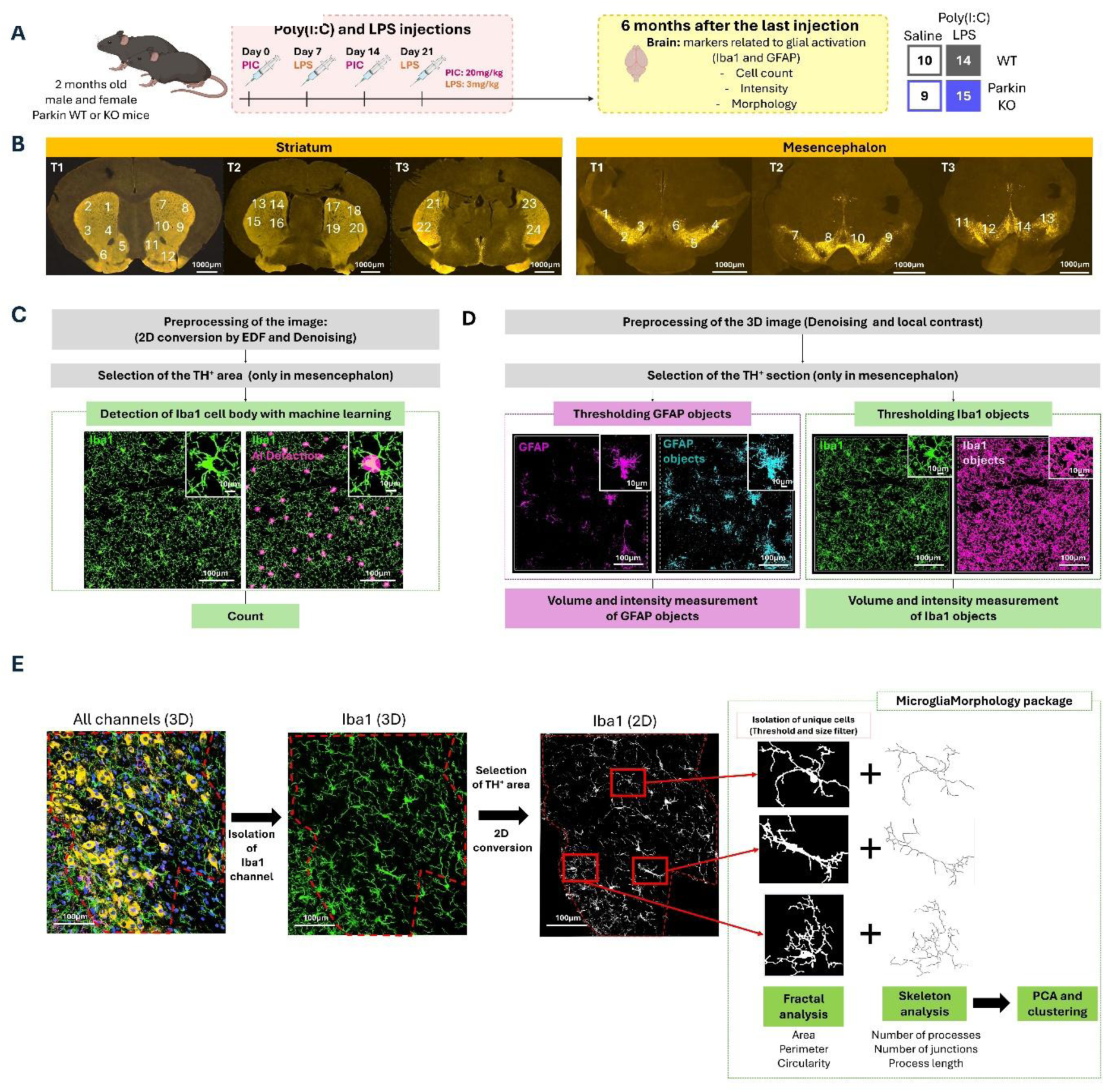
Schematic illustration of the method for semi-automatic analysis of microglia and reactive astrocytes. **(A)** Schematic illustration of the experimental design. WT and Parkin KO mice received alternate Poly(I:C)/LPS injections. Glial activation was evaluated 6 months after the last injection in both striatum and mesencephalon. The number of microglia was counted, the frequency of four microglial morphologies was analysed, and the intensity of Iba1 and GFAP were measured. **(B)** Representative images of immunohistofluorescence showing expression of TH (orange), in three striatal and three mesencephalic slices selected for the study. Images were acquired with a confocal microscope (large image at X20, scale bar: 1000 µm). The numbers represent the regions where the analysed images were quantified to detect microglia and reactive astrocytes. **(C)** Pipeline of semi-automatic microglia counting using the NIS software. After preprocessing of the image, microglia (Iba1, green) and in particular their cells bodies were detected (pink) and counted using machine learning. **(D)** Pipeline of semi-automatic analysis of volume and intensity of Iba1 and GFAP, using the NIS software. After preprocessing of the image, GFAP (magenta) and Iba1 (green) signals were used to create objects (respectively in cyan blue and magenta). The volume and the intensity of the signal were measured in these objects. **(E)** After a 2D reconstruction of the Iba1 signal, the morphology of the microglia was analysed using MicrogliaMophology package on ImageJ. This tool isolates individual cells using intensity and size filters. It then creates masks of these cells and their skeletons. These new images are analyzed to extract numerous parameters and thus cluster the cells into four morphological groups. **C-E**, in mesencephalic slices, the TH area or volume was selected to exclude areas outside of the mesencephalon by setting intensity thresholds, determined based on several test images.

**Supplementary Figure 2:**
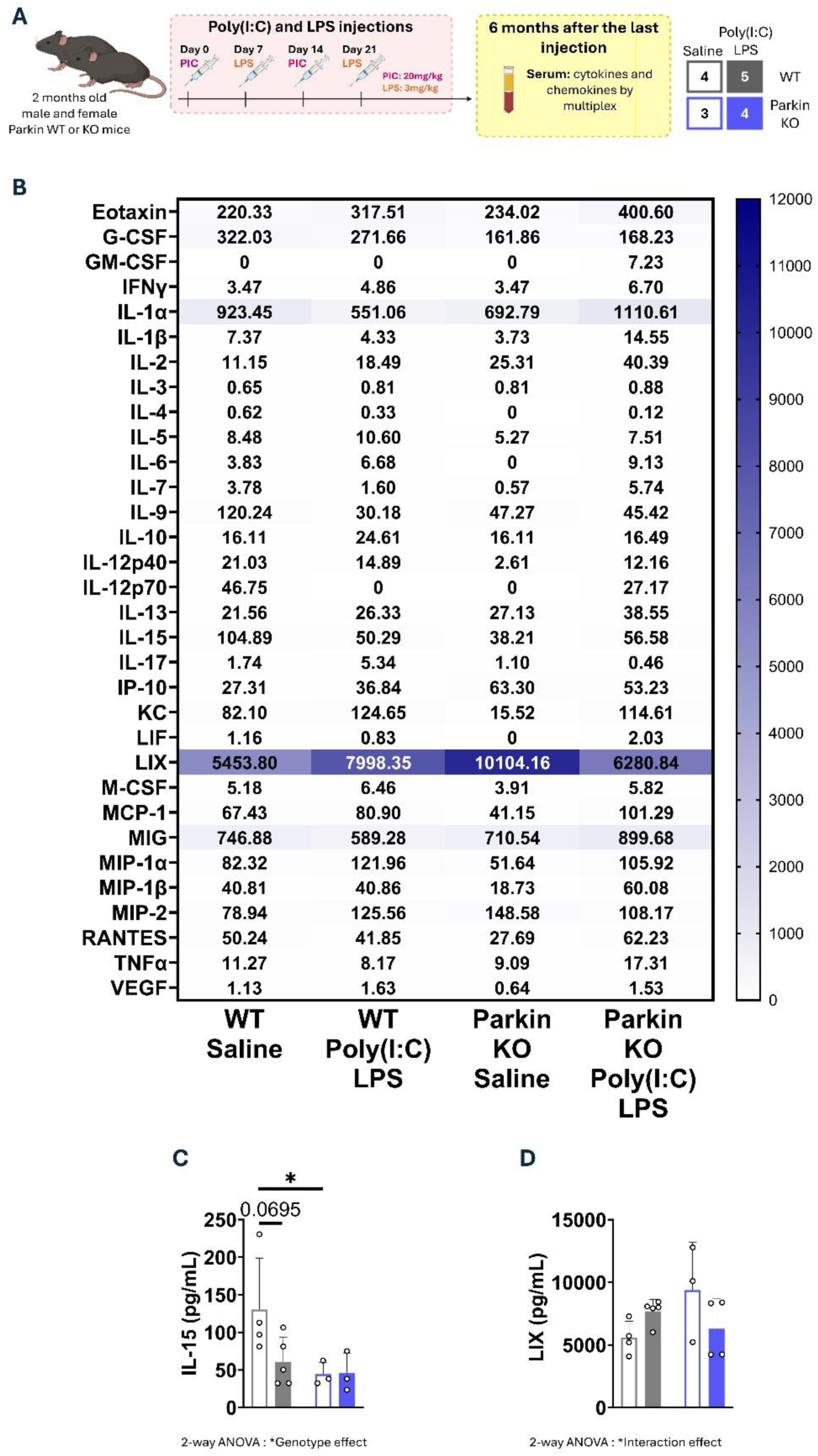
Absence of change in cytokines or chemokines in blood serum 6 months after combined Poly(I:C)/LPS treatment. **(A)** Schematic illustration of experimental design. WT and Parkin KO mice received alternate Poly(I:C)/LPS. Peripheral inflammation was assessed 6 months after the last injection by measuring 32 cytokines and chemokines in serum by a multiplex assay. **(B)** Heatmap showing the concentrations of each cytokine or chemokine. Values indicate group medians, and color intensity follows a continuous gradient, with white representing 0 pg/mL and dark blue representing 12000 pg/mL. **(C)** The serum concentration of IL-15 was lower in Parkin KO mice, and particularly different from their WT counterpart in basal conditions. **(D)** The serum concentration of LIX (CXCL5) was modulated as revealed by an interaction between genotype and treatment. **C-D,** bars represent the mean ± SD of the group and dots represent individual animals. Grey and blue are used respectively for WT and Parkin KO mice. Bars corresponding to saline are unfilled and outlined, while bars corresponding to Poly(I:C)/LPS are filled. Statistical significance was assessed using a 2-way ANOVA on log transformed data (* p<=0.05), the results of which are presented in **Supplementary Table 8**. It was followed by Šídák’s multiple comparisons test (* adjusted p<=0.05) and significant comparisons are indicated above the bars (* adjusted p<=0.05). n=4 for WT Saline, n=5 for WT Poly(I:C)/LPS, n=3 for Parkin KO Saline and n=4 for Parkin KO Poly(I:C)/LPS, from the first cohort only.

**Supplementary Figure 3:**
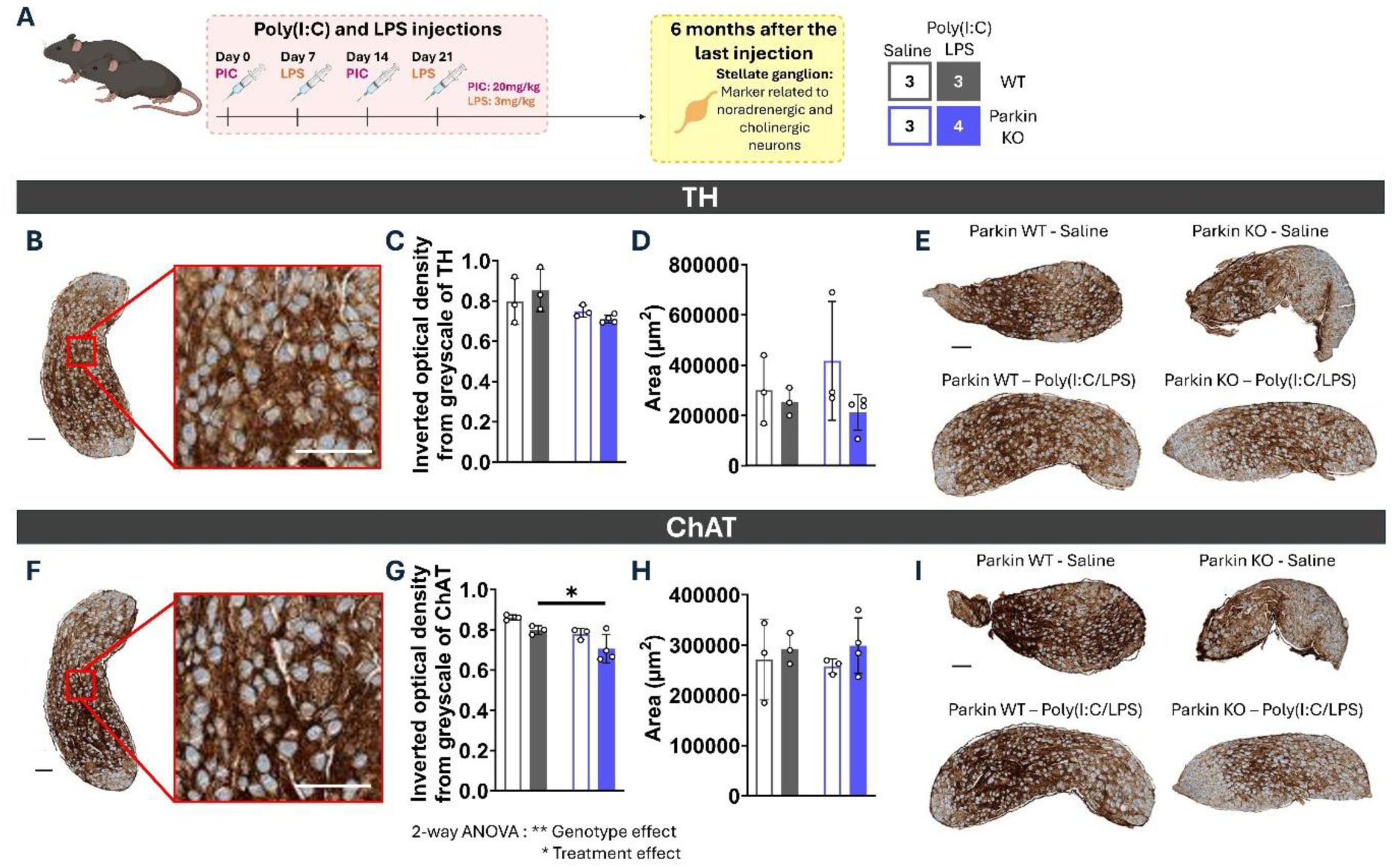
Stellate ganglia show similar TH expression but higher ChAT intensity 6 months after combined Poly(I:C)/LPS treatment in Parkin KO mice. **(A)** Schematic illustration of the experimental design. WT and Parkin KO mice received alternate Poly(I:C)/LPS injections. Peripheral noradrenergic and cholinergic neuronal structures were examined by analysing respectively TH and ChAT markers in stellate ganglia. **(B)** Representative image of TH immunohistochemistry in whole slices of the stellate ganglion (scale bar: 100 µm) and in a zoom of this image (scale bar: 100 µm) showing the TH signal (brown) and the nuclei (blue). **(C)** TH intensity represented by the inverted optical density from greyscale and **(D)** the area covered by the TH signal were similar among the groups. **(E)** Representative image of TH immunohistochemistry in whole slices of the stellate ganglion (scale bar: 100 µm) from Parkin WT and KO mice, treated with saline of Poly(IC)/LPS. TH signal is in brown, and the nuclei is in blue. **(F)** Representative image of ChAT immunohistochemistry in whole slices of the stellate ganglion (scale bar: 100 µm) and in a zoom of this image (scale bar: 100 µm) showing the ChAT signal (brown) and the nuclei (blue). **(G)** ChAT intensity represented by the inverted optical density from greyscale was lower in Parkin KO mice and particularly in mice receiving Poly(I:C)/LPS injections compared to their WT littermates. Statistical significance was assessed using a 2-way ANOVA on log transformed data (* p<=0.05, ** p<0.01), the results of which are presented in **Supplementary Table 10**. It was followed by Šídák’s multiple comparisons test (* adjusted p<=0.05) and significant comparisons are indicated above the bars (* adjusted p<=0.05). **(H)** The area covered by the ChAT signal was comparable between the groups. **(I)** Representative image of ChAT immunohistochemistry in whole slices of the stellate ganglion (scale bar: 100 µm) from Parkin WT and KO mice, treated with saline of Poly(IC)/LPS. ChAT signal is brown and the nuclei is blue. **C-F,** bars represent the mean ± SD of the groups and dots represent individual animals. Grey and blue are used respectively for WT and Parkin KO mice. Bars corresponding to saline are unfilled and outlined, while bars corresponding to Poly(I:C)/LPS are filled. n=3 for WT Saline, n=3 for WT Poly(I:C)/LPS, n=3 for Parkin KO Saline and n=4 for Parkin KO Poly(I:C)/LPS, from the first cohort only.

**Supplementary Table 1:**
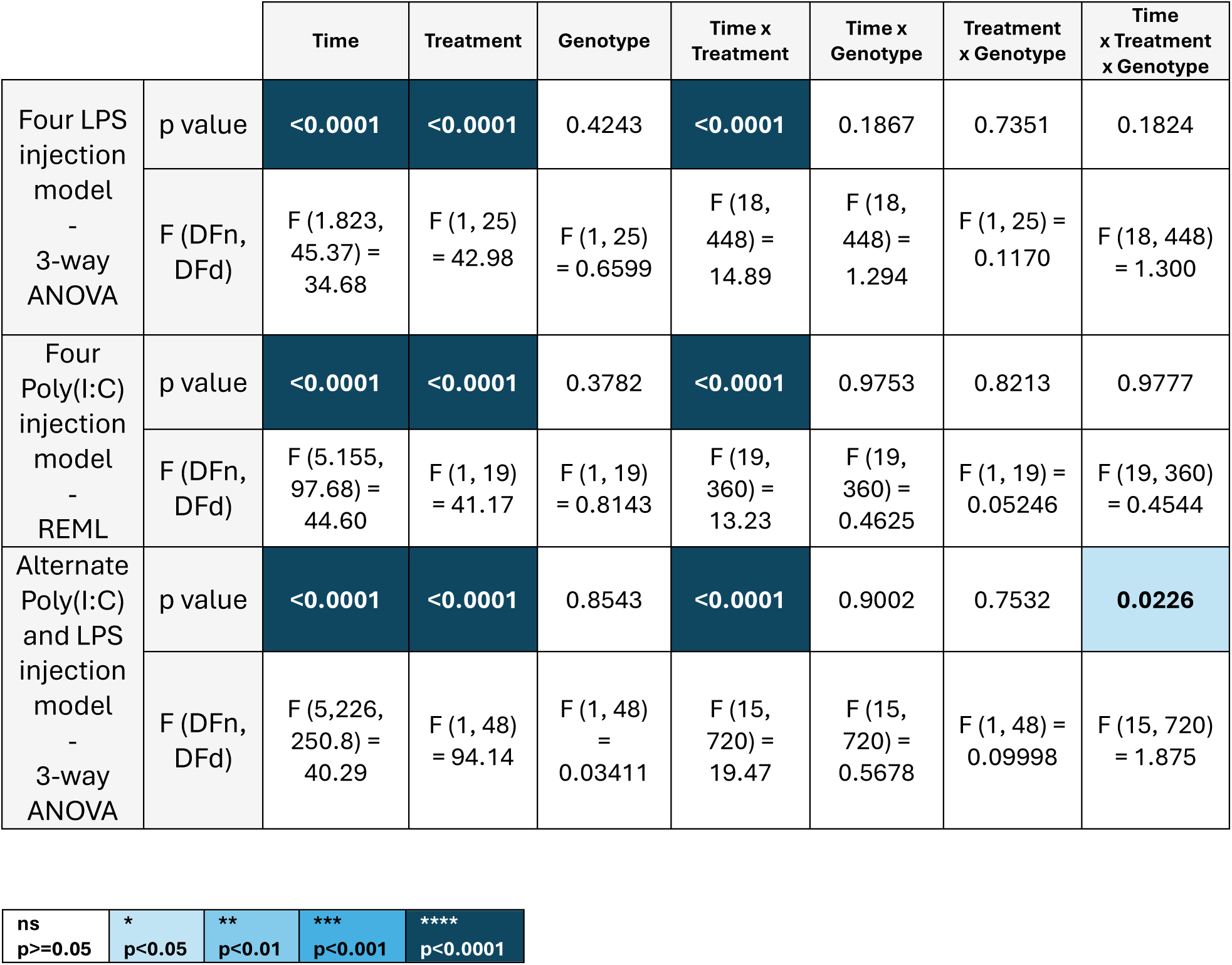
Statistical analysis of the global body weight loss in the three models.

|  |  | Time | Treatment | Genotype | Time x Treatment | Time x Genotype | Treatment x Genotype | Time x Treatment x Genotype |
| --- | --- | --- | --- | --- | --- | --- | --- | --- |
| Four LPS injection model - 3-way ANOVA | p value | <0.0001 | <0.0001 | 0.4243 | <0.0001 | 0.1867 | 0.7351 | 0.1824 |
|  | F (DFn, DFd) | F (1.823, 45.37) = 34.68 | F (1, 25) = 42.98 | F (1, 25) = 0.6599 | F (18, 448) = 14.89 | F (18, 448) = 1.294 | F (1, 25) = 0.1170 | F (18, 448) = 1.300 |
| Four Poly(I:C) injection model - REML | p value | <0.0001 | <0.0001 | 0.3782 | <0.0001 | 0.9753 | 0.8213 | 0.9777 |
|  | F (DFn, DFd) | F (5.155, 97.68) = 44.60 | F (1, 19) = 41.17 | F (1, 19) = 0.8143 | F (19, 360) = 13.23 | F (19, 360) = 0.4625 | F (1, 19) = 0.05246 | F (19, 360) = 0.4544 |
| Alternate Poly(I:C) and LPS injection model - 3-way ANOVA | p value | <0.0001 | <0.0001 | 0.8543 | <0.0001 | 0.9002 | 0.7532 | 0.0226 |
|  | F (DFn, DFd) | F (5,226, 250.8) = 40.29 | F (1, 48) = 94.14 | F (1, 48) = 0.03411 | F (15, 720) = 19.47 | F (15, 720) = 0.5678 | F (1, 48) = 0.09998 | F (15, 720) = 1.875 |

|  |  |  |  |  |
| --- | --- | --- | --- | --- |
| ns<br>p>=0.05 | *<br>p<0.05 | **<br>p<0.01 | ***<br>p<0.001 | ****<br>p<0.0001 |

**Supplementary Table 2:**
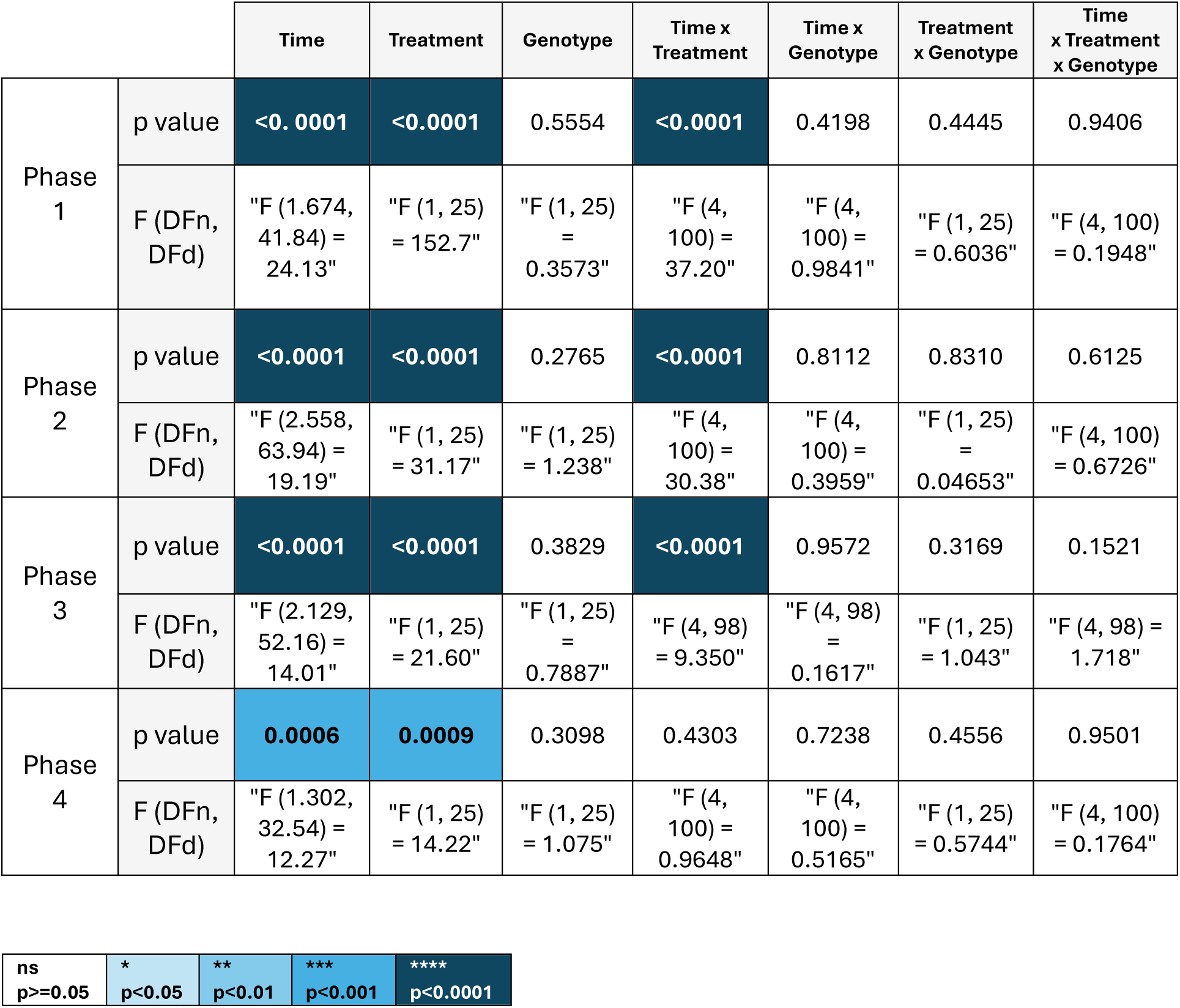
3-way ANOVA of the body weight loss among the different phases in the four LPS injection model.

|  |  | Time | Treatment | Genotype | Time x Treatment | Time x Genotype | Treatment x Genotype | Time x Treatment x Genotype |
| --- | --- | --- | --- | --- | --- | --- | --- | --- |
| Phase 1 | p value | <0.0001 | <0.0001 | 0.5554 | <0.0001 | 0.4198 | 0.4445 | 0.9406 |
|  | F (DFn, DFd) | "F (1.674, 41.84) = 24.13" | "F (1, 25) = 152.7" | "F (1, 25) = 0.3573" | "F (4, 100) = 37.20" | "F (4, 100) = 0.9841" | "F (1, 25) = 0.6036" | "F (4, 100) = 0.1948" |
| Phase 2 | p value | <0.0001 | <0.0001 | 0.2765 | <0.0001 | 0.8112 | 0.8310 | 0.6125 |
|  | F (DFn, DFd) | "F (2.558, 63.94) = 19.19" | "F (1, 25) = 31.17" | "F (1, 25) = 1.238" | "F (4, 100) = 30.38" | "F (4, 100) = 0.3959" | "F (1, 25) = 0.04653" | "F (4, 100) = 0.6726" |
| Phase 3 | p value | <0.0001 | <0.0001 | 0.3829 | <0.0001 | 0.9572 | 0.3169 | 0.1521 |
|  | F (DFn, DFd) | "F (2.129, 52.16) = 14.01" | "F (1, 25) = 21.60" | "F (1, 25) = 0.7887" | "F (4, 98) = 9.350" | "F (4, 98) = 0.1617" | "F (1, 25) = 1.043" | "F (4, 98) = 1.718" |
| Phase 4 | p value | 0.0006 | 0.0009 | 0.3098 | 0.4303 | 0.7238 | 0.4556 | 0.9501 |
|  | F (DFn, DFd) | "F (1.302, 32.54) = 12.27" | "F (1, 25) = 14.22" | "F (1, 25) = 1.075" | "F (4, 100) = 0.9648" | "F (4, 100) = 0.5165" | "F (1, 25) = 0.5744" | "F (4, 100) = 0.1764" |

|  |  |  |  |  |
| --- | --- | --- | --- | --- |
| ns<br>p>=0.05 | *<br>p<0.05 | **<br>p<0.01 | ***<br>p<0.001 | ****<br>p<0.0001 |

**Supplementary Table 3:**
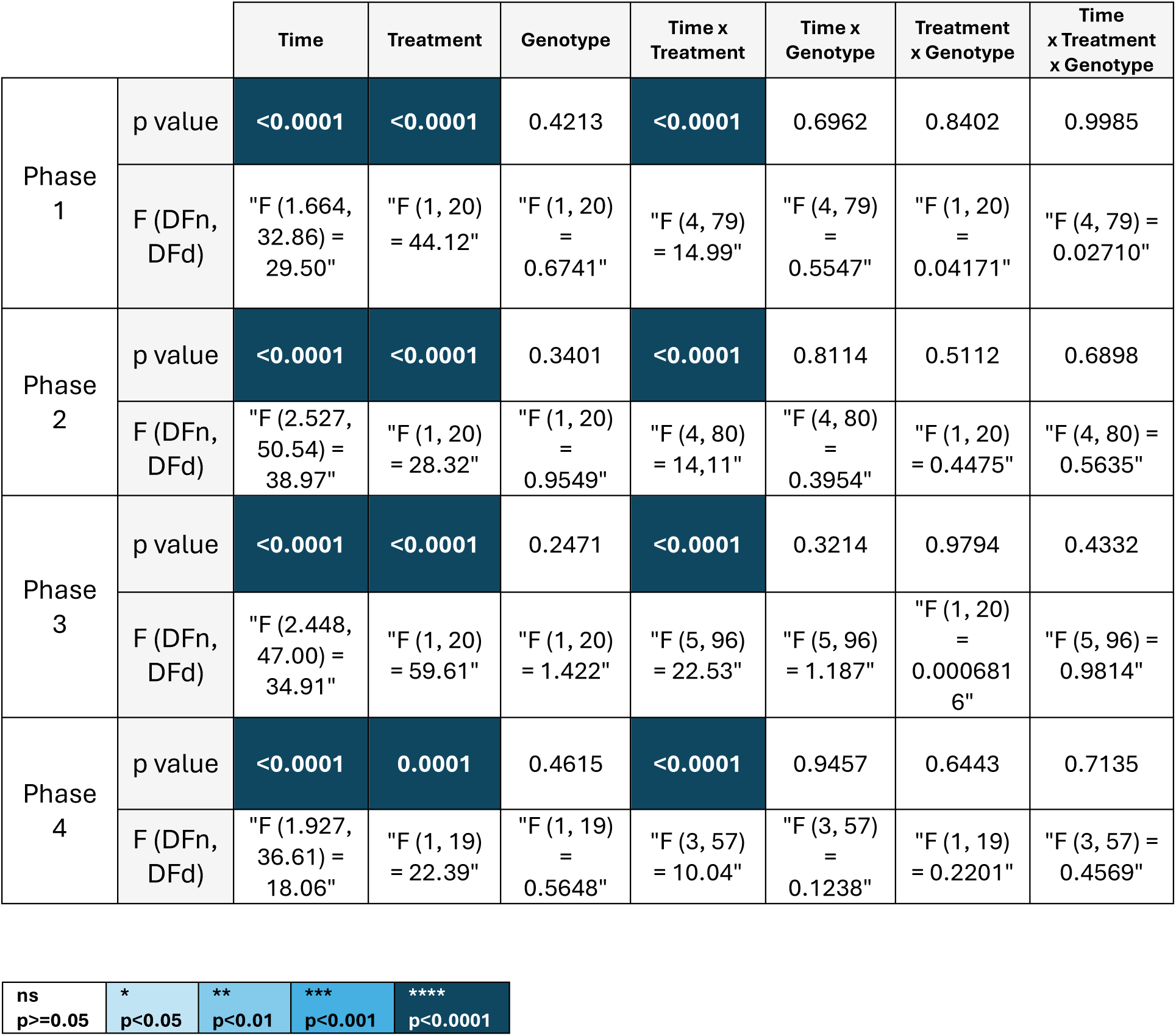
3-way ANOVA of the body weight loss among the different phases in the four Poly(I:C) injection model.

|  |  | Time | Treatment | Genotype | Time x Treatment | Time x Genotype | Treatment x Genotype | Time x Treatment x Genotype |
| --- | --- | --- | --- | --- | --- | --- | --- | --- |
| Phase 1 | p value | <0.0001 | <0.0001 | 0.4213 | <0.0001 | 0.6962 | 0.8402 | 0.9985 |
|  | F (DFn, DFd) | "F (1.664, 32.86) = 29.50" | "F (1, 20) = 44.12" | "F (1, 20) = 0.6741" | "F (4, 79) = 14.99" | "F (4, 79) = 0.5547" | "F (1, 20) = 0.04171" | "F (4, 79) = 0.02710" |
| Phase 2 | p value | <0.0001 | <0.0001 | 0.3401 | <0.0001 | 0.8114 | 0.5112 | 0.6898 |
|  | F (DFn, DFd) | "F (2.527, 50.54) = 38.97" | "F (1, 20) = 28.32" | "F (1, 20) = 0.9549" | "F (4, 80) = 14,11" | "F (4, 80) = 0.3954" | "F (1, 20) = 0.4475" | "F (4, 80) = 0.5635" |
| Phase 3 | p value | <0.0001 | <0.0001 | 0.2471 | <0.0001 | 0.3214 | 0.9794 | 0.4332 |
|  | F (DFn, DFd) | "F (2.448, 47.00) = 34.91" | "F (1, 20) = 59.61" | "F (1, 20) = 1.422" | "F (5, 96) = 22.53" | "F (5, 96) = 1.187" | "F (1, 20) = 0.0006816" | "F (5, 96) = 0.9814" |
| Phase 4 | p value | <0.0001 | 0.0001 | 0.4615 | <0.0001 | 0.9457 | 0.6443 | 0.7135 |
|  | F (DFn, DFd) | "F (1.927, 36.61) = 18.06" | "F (1, 19) = 22.39" | "F (1, 19) = 0.5648" | "F (3, 57) = 10.04" | "F (3, 57) = 0.1238" | "F (1, 19) = 0.2201" | "F (3, 57) = 0.4569" |

|  |  |  |  |  |
| --- | --- | --- | --- | --- |
| ns<br>p>=0.05 | *<br>p<0.05 | **<br>p<0.01 | ***<br>p<0.001 | ****<br>p<0.0001 |

**Supplementary Table 4:**
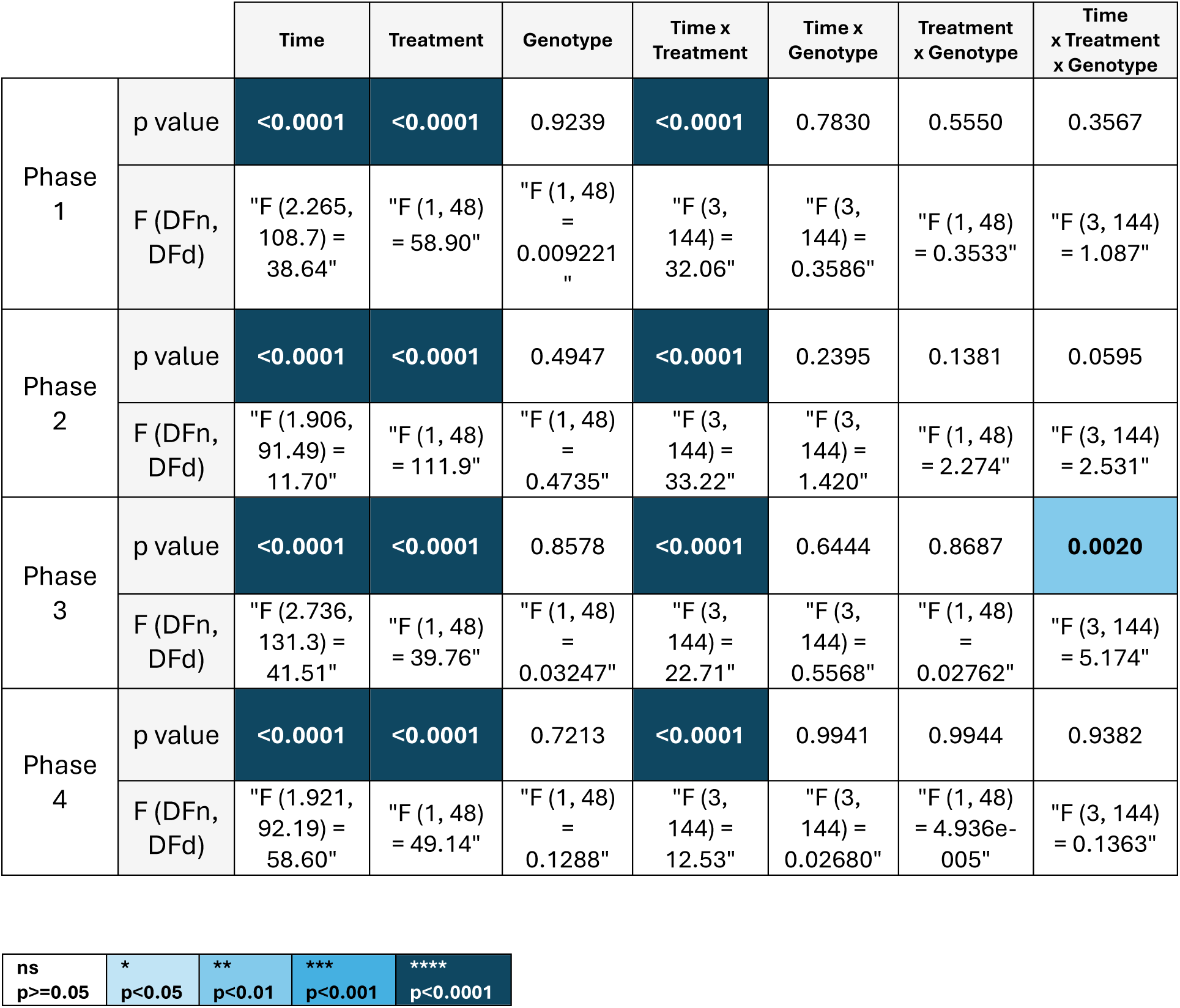
3-way ANOVA of the body weight loss among the different phases in the alternate Poly(I:C)/LPS injection model.

|  |  | Time | Treatment | Genotype | Time x Treatment | Time x Genotype | Treatment x Genotype | Time x Treatment x Genotype |
| --- | --- | --- | --- | --- | --- | --- | --- | --- |
| Phase 1 | p value | <0.0001 | <0.0001 | 0.9239 | <0.0001 | 0.7830 | 0.5550 | 0.3567 |
|  | F (DFn, DFd) | "F (2.265, 108.7) = 38.64" | "F (1, 48) = 58.90" | "F (1, 48) = 0.009221" | "F (3, 144) = 32.06" | "F (3, 144) = 0.3586" | "F (1, 48) = 0.3533" | "F (3, 144) = 1.087" |
| Phase 2 | p value | <0.0001 | <0.0001 | 0.4947 | <0.0001 | 0.2395 | 0.1381 | 0.0595 |
|  | F (DFn, DFd) | "F (1.906, 91.49) = 11.70" | "F (1, 48) = 111.9" | "F (1, 48) = 0.4735" | "F (3, 144) = 33.22" | "F (3, 144) = 1.420" | "F (1, 48) = 2.274" | "F (3, 144) = 2.531" |
| Phase 3 | p value | <0.0001 | <0.0001 | 0.8578 | <0.0001 | 0.6444 | 0.8687 | 0.0020 |
|  | F (DFn, DFd) | "F (2.736, 131.3) = 41.51" | "F (1, 48) = 39.76" | "F (1, 48) = 0.03247" | "F (3, 144) = 22.71" | "F (3, 144) = 0.5568" | "F (1, 48) = 0.02762" | "F (3, 144) = 5.174" |
| Phase 4 | p value | <0.0001 | <0.0001 | 0.7213 | <0.0001 | 0.9941 | 0.9944 | 0.9382 |
|  | F (DFn, DFd) | "F (1.921, 92.19) = 58.60" | "F (1, 48) = 49.14" | "F (1, 48) = 0.1288" | "F (3, 144) = 12.53" | "F (3, 144) = 0.02680" | "F (1, 48) = 4.936e-005" | "F (3, 144) = 0.1363" |

|  |  |  |  |  |
| --- | --- | --- | --- | --- |
| ns<br>p>=0.05 | *<br>p<0.05 | **<br>p<0.01 | ***<br>p<0.001 | ****<br>p<0.0001 |

**Supplementary Table 5:**
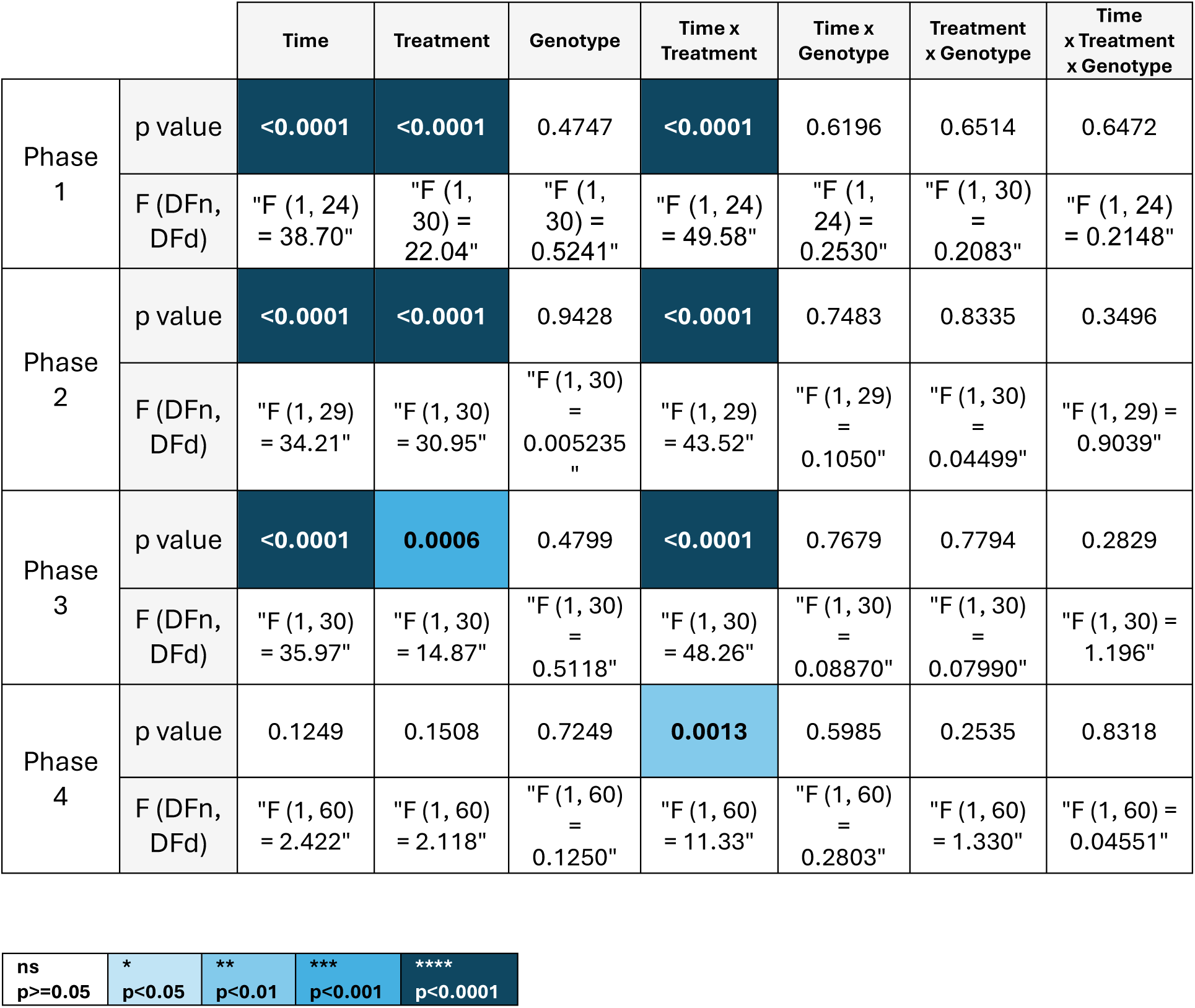
3-way ANOVA of the lipocalin level among the different phases in the alternate Poly(I:C)/LPS injection model.

|  |  | Time | Treatment | Genotype | Time x Treatment | Time x Genotype | Treatment x Genotype | Time x Treatment x Genotype |
| --- | --- | --- | --- | --- | --- | --- | --- | --- |
| Phase 1 | p value | <0.0001 | <0.0001 | 0.4747 | <0.0001 | 0.6196 | 0.6514 | 0.6472 |
|  | F (DFn, DFd) | "F (1, 24) = 38.70" | "F (1, 30) = 22.04" | "F (1, 30) = 0.5241" | "F (1, 24) = 49.58" | "F (1, 24) = 0.2530" | "F (1, 30) = 0.2083" | "F (1, 24) = 0.2148" |
| Phase 2 | p value | <0.0001 | <0.0001 | 0.9428 | <0.0001 | 0.7483 | 0.8335 | 0.3496 |
|  | F (DFn, DFd) | "F (1, 29) = 34.21" | "F (1, 30) = 30.95" | "F (1, 30) = 0.005235" | "F (1, 29) = 43.52" | "F (1, 29) = 0.1050" | "F (1, 30) = 0.04499" | "F (1, 29) = 0.9039" |
| Phase 3 | p value | <0.0001 | 0.0006 | 0.4799 | <0.0001 | 0.7679 | 0.7794 | 0.2829 |
|  | F (DFn, DFd) | "F (1, 30) = 35.97" | "F (1, 30) = 14.87" | "F (1, 30) = 0.5118" | "F (1, 30) = 48.26" | "F (1, 30) = 0.08870" | "F (1, 30) = 0.07990" | "F (1, 30) = 1.196" |
| Phase 4 | p value | 0.1249 | 0.1508 | 0.7249 | 0.0013 | 0.5985 | 0.2535 | 0.8318 |
|  | F (DFn, DFd) | "F (1, 60) = 2.422" | "F (1, 60) = 2.118" | "F (1, 60) = 0.1250" | "F (1, 60) = 11.33" | "F (1, 60) = 0.2803" | "F (1, 60) = 1.330" | "F (1, 60) = 0.04551" |

|  |  |  |  |  |
| --- | --- | --- | --- | --- |
| ns<br>p>=0.05 | *<br>p<0.05 | **<br>p<0.01 | ***<br>p<0.001 | ****<br>p<0.0001 |

**Supplementary Table 6:**
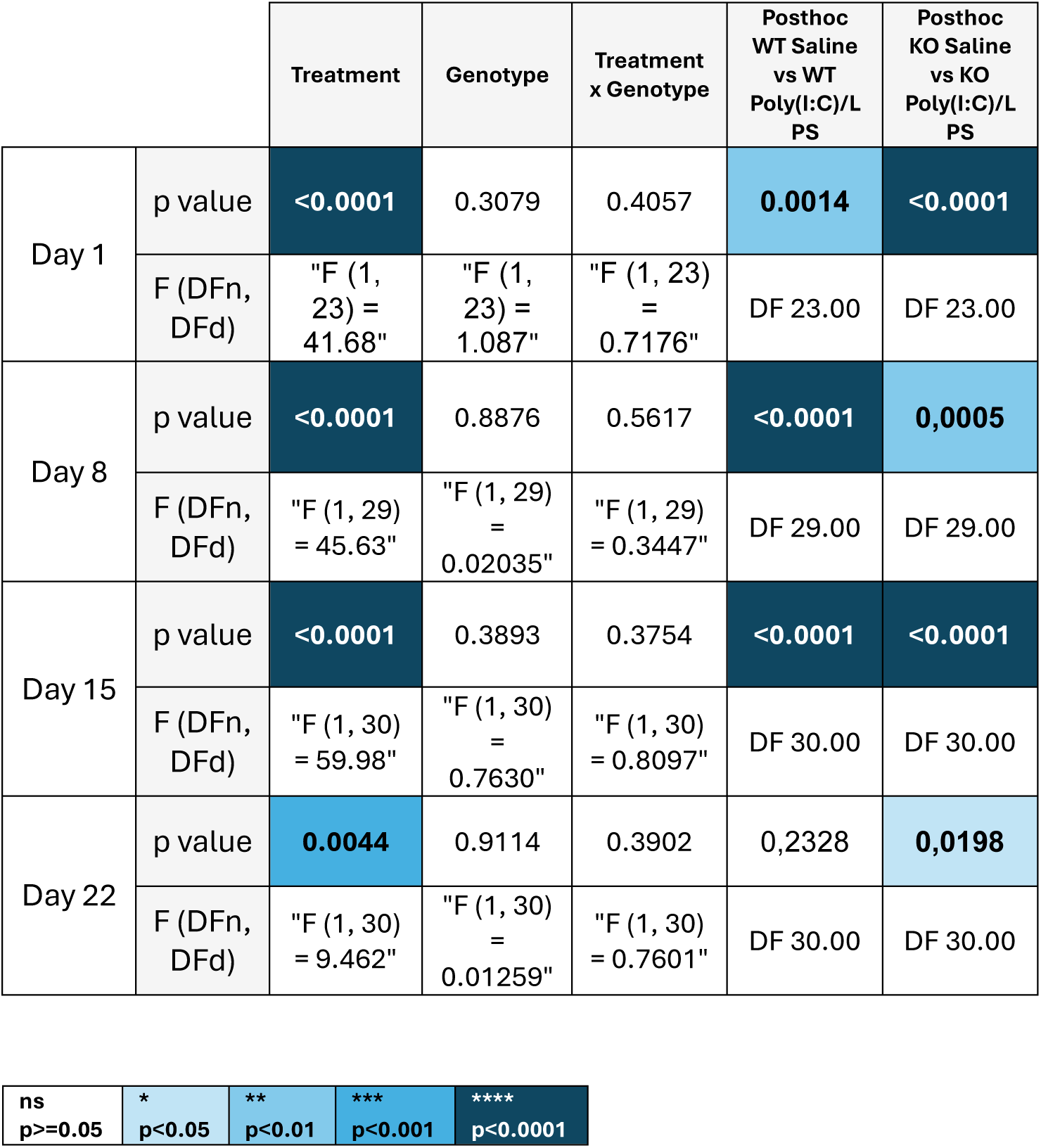
2-way ANOVA of the fecal lipocalin level each day after injections in the alternate Poly(I:C)/LPS model.

|  |  | Treatment | Genotype | Treatment<br>x Genotype | Posthoc<br>WT Saline<br>vs WT<br>Poly(I:C)/L<br>PS | Posthoc<br>KO Saline<br>vs KO<br>Poly(I:C)/L<br>PS |
| --- | --- | --- | --- | --- | --- | --- |
| Day 1 | p value | <0.0001 | 0.3079 | 0.4057 | 0.0014 | <0.0001 |
|  | F (DFn, DFd) | "F (1, 23) = 41.68" | "F (1, 23) = 1.087" | "F (1, 23) = 0.7176" | DF 23.00 | DF 23.00 |
| Day 8 | p value | <0.0001 | 0.8876 | 0.5617 | <0.0001 | 0,0005 |
|  | F (DFn, DFd) | "F (1, 29) = 45.63" | "F (1, 29) = 0.02035" | "F (1, 29) = 0.3447" | DF 29.00 | DF 29.00 |
| Day 15 | p value | <0.0001 | 0.3893 | 0.3754 | <0.0001 | <0.0001 |
|  | F (DFn, DFd) | "F (1, 30) = 59.98" | "F (1, 30) = 0.7630" | "F (1, 30) = 0.8097" | DF 30.00 | DF 30.00 |
| Day 22 | p value | 0.0044 | 0.9114 | 0.3902 | 0,2328 | 0,0198 |
|  | F (DFn, DFd) | "F (1, 30) = 9.462" | "F (1, 30) = 0.01259" | "F (1, 30) = 0.7601" | DF 30.00 | DF 30.00 |

|  |  |  |  |  |
| --- | --- | --- | --- | --- |
| ns<br>p>=0.05 | *<br>p<0.05 | **<br>p<0.01 | ***<br>p<0.001 | ****<br>p<0.0001 |

**Supplementary Table 7:**
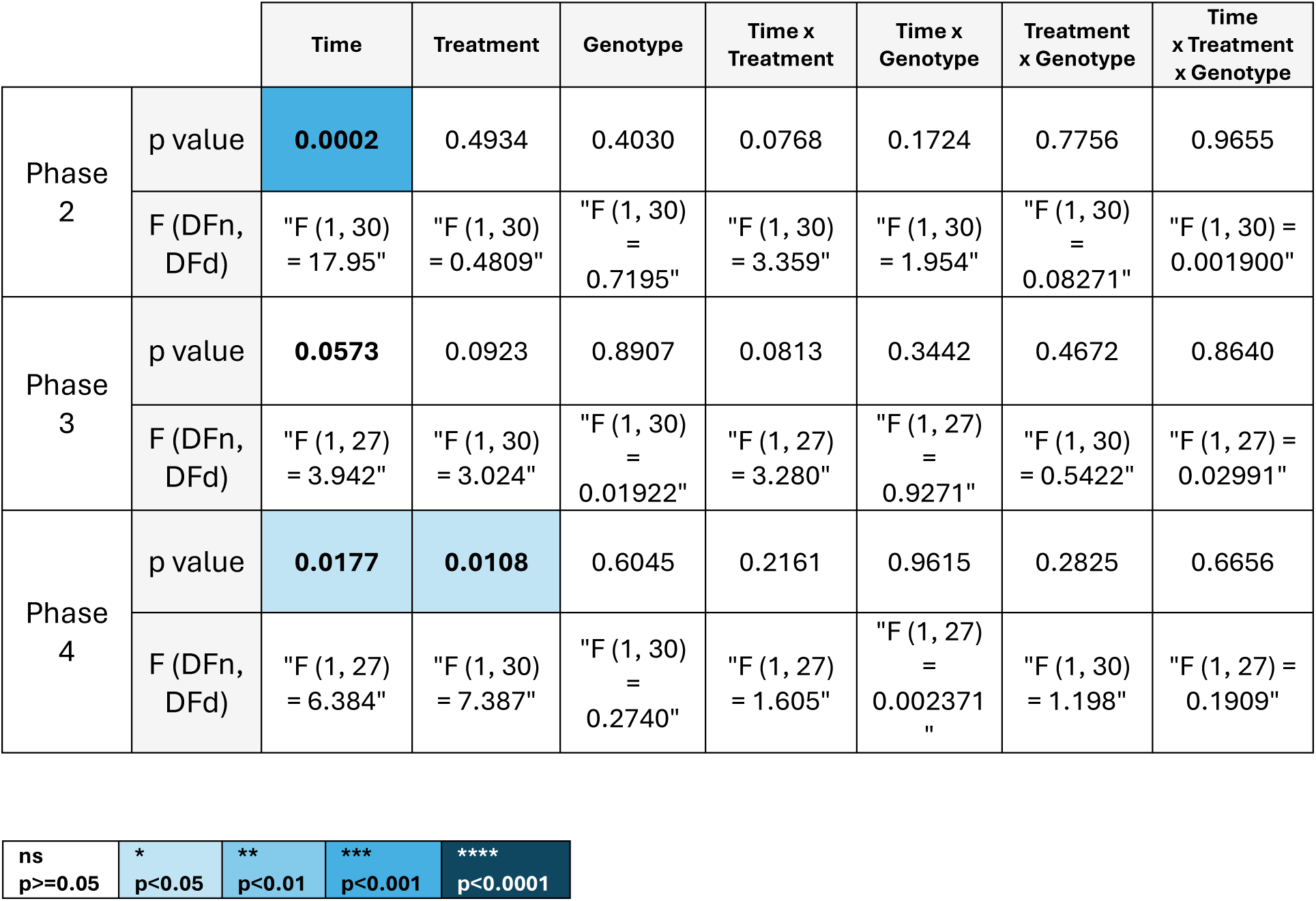
3-way ANOVA of the calprotectin level among the different phases in the alternate Poly(I:C)/LPS injection model.

|  |  | Time | Treatment | Genotype | Time x Treatment | Time x Genotype | Treatment x Genotype | Time x Treatment x Genotype |
| --- | --- | --- | --- | --- | --- | --- | --- | --- |
| Phase 2 | p value | 0.0002 | 0.4934 | 0.4030 | 0.0768 | 0.1724 | 0.7756 | 0.9655 |
|  | F (DFn, DFd) | "F (1, 30) = 17.95" | "F (1, 30) = 0.4809" | "F (1, 30) = 0.7195" | "F (1, 30) = 3.359" | "F (1, 30) = 1.954" | "F (1, 30) = 0.08271" | "F (1, 30) = 0.001900" |
| Phase 3 | p value | 0.0573 | 0.0923 | 0.8907 | 0.0813 | 0.3442 | 0.4672 | 0.8640 |
|  | F (DFn, DFd) | "F (1, 27) = 3.942" | "F (1, 30) = 3.024" | "F (1, 30) = 0.01922" | "F (1, 27) = 3.280" | "F (1, 27) = 0.9271" | "F (1, 30) = 0.5422" | "F (1, 27) = 0.02991" |
| Phase 4 | p value | 0.0177 | 0.0108 | 0.6045 | 0.2161 | 0.9615 | 0.2825 | 0.6656 |
|  | F (DFn, DFd) | "F (1, 27) = 6.384" | "F (1, 30) = 7.387" | "F (1, 30) = 0.2740" | "F (1, 27) = 1.605" | "F (1, 27) = 0.002371" | "F (1, 30) = 1.198" | "F (1, 27) = 0.1909" |

|  |  |  |  |  |
| --- | --- | --- | --- | --- |
| ns<br>p>=0.05 | *<br>p<0.05 | **<br>p<0.01 | ***<br>p<0.001 | ****<br>p<0.0001 |

**Supplementary Table 8:** 2-way ANOVA of the fecal calprotectin level, the spleen index, the levels of IL-15 and LIX six months after the alternate Poly(I:C)/LPS injection.

|  |  | Treatment | Genotype | Treatment<br>x Genotype |
| --- | --- | --- | --- | --- |
| Fecal<br>calprot<br>ectin | p value | 0.7894 | **0.0378** | 0.1348 |
|  | F (DFn,<br>DFd) | "F (1, 46)<br>=<br>0.07215" | "F (1, 46)<br>= 4.575" | "F (1, 46)<br>= 2.317" |
| Spleen<br>index | p value | 0.4420 | **0.0323** | 0.3901 |
|  | F (DFn,<br>DFd) | "F (1, 44)<br>= 0.6018" | "F (1, 44)<br>= 4.886" | "F (1, 44)<br>= 0.7533" |
| Level<br>of IL-15 | p value | 0.1356 | **0.0292** | 0.1794 |
|  | F (DFn,<br>DFd) | "F (1, 11)<br>= 2.593" | "F (1, 11)<br>= 6.283" | "F (1, 11)<br>= 2.057" |
| Level<br>of LIX | p value | 0.6682 | 0.2917 | **0.0347** |
|  | F (DFn,<br>DFd) | "F (1, 12)<br>= 0.1931" | "F (1, 12)<br>= 1.217" | "F (1, 12)<br>= 5.672" |
| ns<br>p>=0.05 | *<br>p<0.05 | **<br>p<0.01 | ***<br>p<0.001 | ****<br>p<0.0001 |

**Supplementary Table 9:**
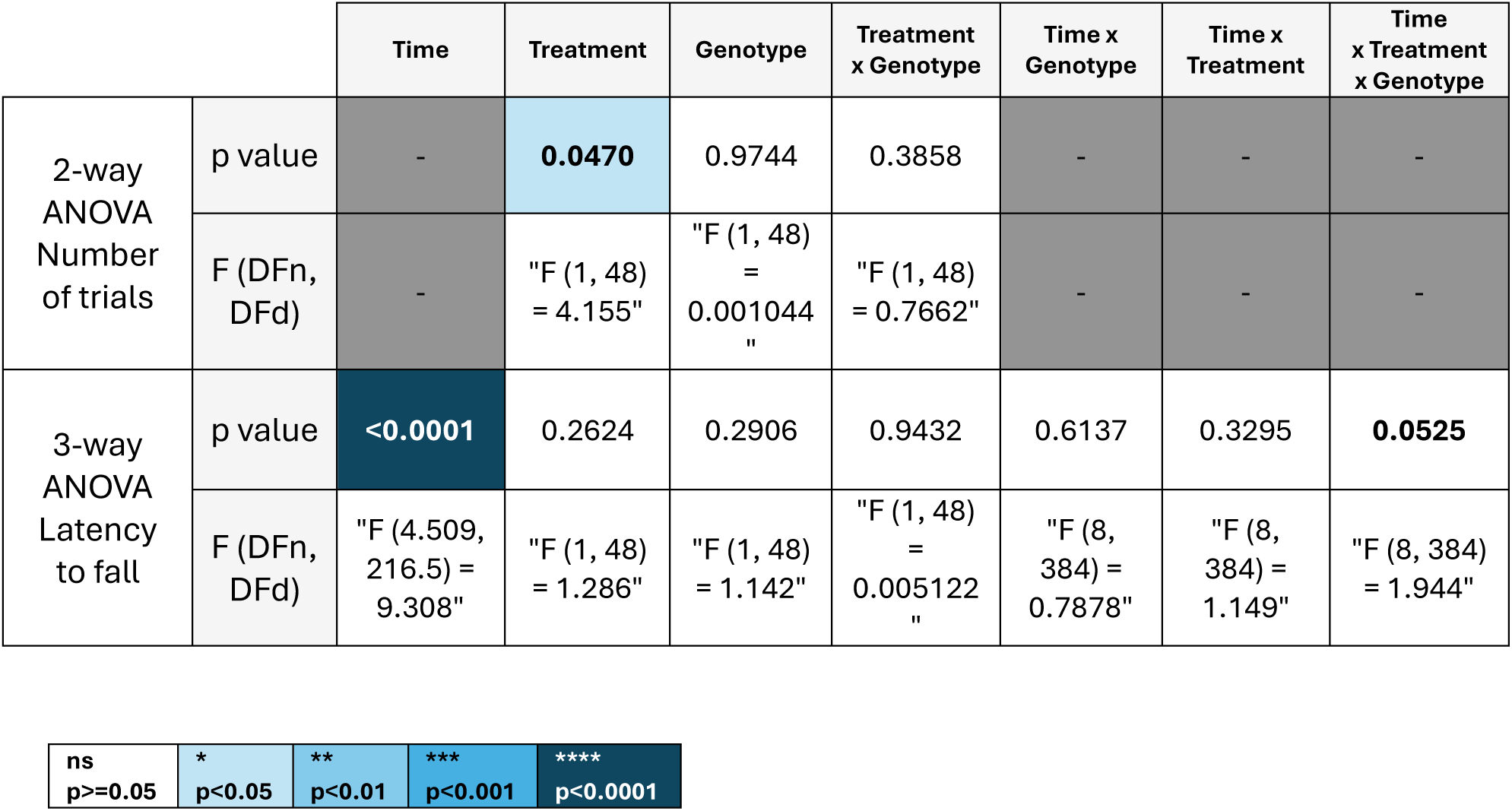
2-way and 3-way ANOVA of the rotarod four months after the alternate Poly(I:C)/LPS injection.

|  |  | Time | Treatment | Genotype | Treatment<br>x Genotype | Time x<br>Genotype | Time x<br>Treatment | Time<br>x Treatment<br>x Genotype |
| --- | --- | --- | --- | --- | --- | --- | --- | --- |
| 2-way<br>ANOVA<br>Number<br>of trials | p value | - | **0.0470** | 0.9744 | 0.3858 | - | - | - |
|  | F (DFn,<br>DFd) | - | "F (1, 48)<br>= 4.155" | "F (1, 48)<br>=<br>0.001044<br>" | "F (1, 48)<br>=<br>0.7662" | - | - | - |
| 3-way<br>ANOVA<br>Latency<br>to fall | p value | **<0.0001** | 0.2624 | 0.2906 | 0.9432 | 0.6137 | 0.3295 | **0.0525** |
|  | F (DFn,<br>DFd) | "F (4.509,<br>216.5) =<br>9.308" | "F (1, 48)<br>= 1.286" | "F (1, 48)<br>= 1.142" | "F (1, 48)<br>=<br>0.005122<br>" | "F (8,<br>384) =<br>0.7878" | "F (8,<br>384) =<br>1.149" | "F (8, 384)<br>= 1.944" |

|  |  |  |  |  |
| --- | --- | --- | --- | --- |
| ns<br>p>=0.05 | *<br>p<0.05 | **<br>p<0.01 | ***<br>p<0.001 | ****<br>p<0.0001 |

**Supplementary Table 10:** 2-way ANOVA of ChAT intensity in stellate ganglion six months after the alternate Poly(I:C)/LPS injection.

|  |  | Treatment | Genotype | Treatment<br>x Genotype |
| --- | --- | --- | --- | --- |
| CHAT<br>intensity | p value | 0.0217 | 0.0065 | 0.8285 |
|  | F (DFn,<br>DFd) | "F (1, 9) =<br>7.674" | "F (1, 9) =<br>12.44" | "F (1, 9) =<br>0.04975" |
| ns<br>p>=0.05 | *<br>p<0.05 | **<br>p<0.01 | ***<br>p<0.001 | ****<br>p<0.0001 |

**Supplementary Table 11:**
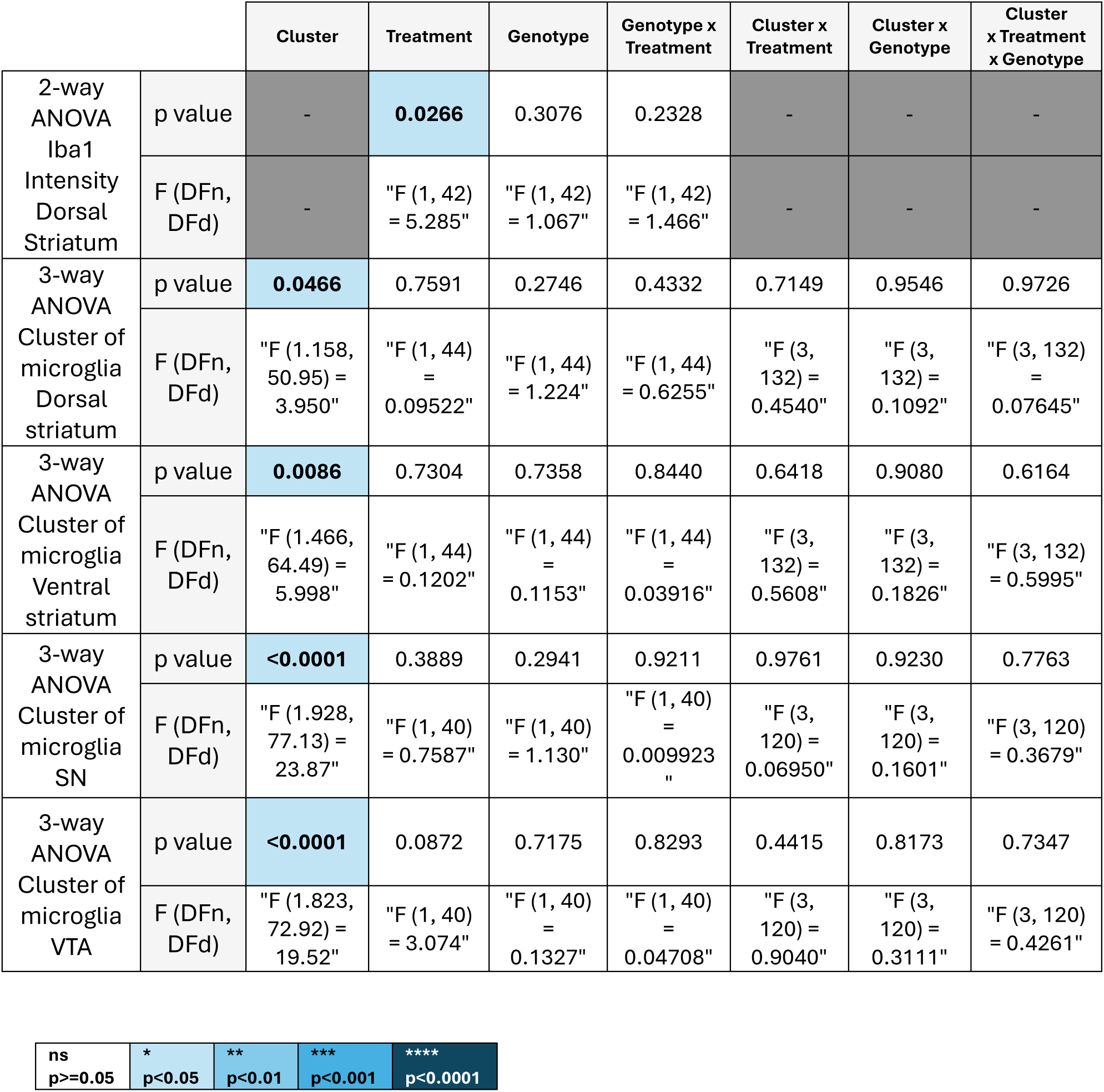
2-way and 3-way ANOVA of microglial analysis six months after the alternate Poly(I:C)/LPS injection.

|  |  | Cluster | Treatment | Genotype | Genotype x Treatment | Cluster x Treatment | Cluster x Genotype | Cluster x Treatment x Genotype |
| --- | --- | --- | --- | --- | --- | --- | --- | --- |
| 2-way ANOVA Iba1 Intensity Dorsal Striatum | p value | - | **0.0266** | 0.3076 | 0.2328 | - | - | - |
|  | F (DFn, DFd) | - | "F (1, 42) = 5.285" | "F (1, 42) = 1.067" | "F (1, 42) = 1.466" | - | - | - |
| 3-way ANOVA Cluster of microglia Dorsal striatum | p value | **0.0466** | 0.7591 | 0.2746 | 0.4332 | 0.7149 | 0.9546 | 0.9726 |
|  | F (DFn, DFd) | "F (1.158, 50.95) = 3.950" | "F (1, 44) = 0.09522" | "F (1, 44) = 1.224" | "F (1, 44) = 0.6255" | "F (3, 132) = 0.4540" | "F (3, 132) = 0.1092" | "F (3, 132) = 0.07645" |
| 3-way ANOVA Cluster of microglia Ventral striatum | p value | **0.0086** | 0.7304 | 0.7358 | 0.8440 | 0.6418 | 0.9080 | 0.6164 |
|  | F (DFn, DFd) | "F (1.466, 64.49) = 5.998" | "F (1, 44) = 0.1202" | "F (1, 44) = 0.1153" | "F (1, 44) = 0.03916" | "F (3, 132) = 0.5608" | "F (3, 132) = 0.1826" | "F (3, 132) = 0.5995" |
| 3-way ANOVA Cluster of microglia SN | p value | **<0.0001** | 0.3889 | 0.2941 | 0.9211 | 0.9761 | 0.9230 | 0.7763 |
|  | F (DFn, DFd) | "F (1.928, 77.13) = 23.87" | "F (1, 40) = 0.7587" | "F (1, 40) = 1.130" | "F (1, 40) = 0.009923" | "F (3, 120) = 0.06950" | "F (3, 120) = 0.1601" | "F (3, 120) = 0.3679" |
| 3-way ANOVA Cluster of microglia VTA | p value | **<0.0001** | 0.0872 | 0.7175 | 0.8293 | 0.4415 | 0.8173 | 0.7347 |
|  | F (DFn, DFd) | "F (1.823, 72.92) = 19.52" | "F (1, 40) = 3.074" | "F (1, 40) = 0.1327" | "F (1, 40) = 0.04708" | "F (3, 120) = 0.9040" | "F (3, 120) = 0.3111" | "F (3, 120) = 0.4261" |

|  |  |  |  |  |
| --- | --- | --- | --- | --- |
| ns | * | ** | *** | **** |
| p>=0.05 | p<0.05 | p<0.01 | p<0.001 | p<0.0001 |

**Supplementary Table 12:** 2-way ANOVA of astrocytes analysis six months after the alternate Poly(I:C)/LPS injection.

|  |  | Treatment | Genotype | Genotype x Treatment |
| --- | --- | --- | --- | --- |
| 2-way ANOVA<br>Count of GFAP cells SN | p value | 0.0341 | 0.4766 | 0.0641 |
|  | F (DFn, DFd) | "F (1, 40) = 4.816" | "F (1, 40) = 0.5164" | "F (1, 40) = 3.627" |
| 2-way ANOVA<br>Percent volume of GFAP SN | p value | 0.1575 | 0.2005 | 0.0117 |
|  | F (DFn, DFd) | "F (1, 40) = 2.075" | "F (1, 40) = 1.694" | "F (1, 40) = 6.975" |
| 2-way ANOVA<br>Count of GFAP cells VTA | p value | 0.0584 | 0.6665 | 0.0273 |
|  | F (DFn, DFd) | "F (1, 40) = 3.797" | "F (1, 40) = 0.1885" | "F (1, 40) = 5.251" |
| 2-way ANOVA GFAP fluorescence VTA | p value | 0.0228 | 0.3241 | 0.8605 |
|  | F (DFn, DFd) | "F (1, 40) = 5.609" | "F (1, 40) = 0.9968" | "F (1, 40) = 0.03129" |
| ns<br>p>=0.05 | *<br>p<0.05 | **<br>p<0.01 | ***<br>p<0.001 | ****<br>p<0.0001 |

